# Complementary chemical and effect-based risk assessments reveal ecotoxicological risks in Caribbean coastal bays and coral reefs

**DOI:** 10.64898/2026.11.02.763482

**Authors:** N.C.J. van de Loosdrecht, M.L. de Baat, M.J.A. Vermeij, P. Aguilar-Alarcón, P.M. Visser, J.M. de Goeij

**Author notes:** corresponding author: N.C.J. van de Loosdrecht.

## Abstract

Anthropogenic chemical pollution poses a significant, yet often unquantified threat to coral reef ecosystems. This study investigated the occurrence, dispersal, and ecotoxicological risks of chemical mixtures in three inland bays of Curaçao (Southern Caribbean) and adjacent coral reefs during wet and dry seasons. Passive sampler extracts were analyzed with targeted chemical analysis and effect-based methods. Inland bays acted as reservoirs for chemical pollution, each exhibiting a unique chemical fingerprint likely linked to surrounding land use. Targeted chemical concentrations were higher during the wet season, particularly inside the bays, indicating rainfall-driven mobilization of terrestrial contaminants. Many compounds exceeded their predicted no-effect concentrations, while mixture toxic pressure was low to moderate and highest inside the bays. In contrast, bioassays integrating the effects of the entire mixture revealed a potential ecotoxicological risk, with activation of the Nrf2 oxidative stress pathway dominating across reef sites. The two lines of evidence were complementary: target compounds identify the bays as pollution sources, while bioassays show mixture effects are strongest on the reefs. Hence, targeted chemical analysis alone is insufficient to evaluate ecotoxicological risk. Our multidisciplinary approach provides a framework for prioritizing key chemical indicators and bioassays to monitor coral reef water quality.

## 1. Introduction

Coral reefs are one of the most biodiverse and productive ecosystems on Earth [1, 2] that deliver crucial ecosystem services to nearby communities, such as fisheries, coastal protection and tourism, valued at hundreds of billions of US dollars per year [3, 4]. However, global climate change [5] and local anthropogenic stressors [6] have jointly contributed to the ongoing degradation of coral reefs around the world. Of these local stressors, overfishing and pollution from land-based human activities are two of the most important for coastal coral reefs [7, 8]. Unsustainable coastal development, intensive agriculture, and industrialization have all contributed to a growing input and higher diversity of pollutants entering nearshore waters [7–9] via terrestrial runoff [10, 11], direct discharge [12–14], and submarine groundwater flow [15]. Pollutants comprise a complex mixture of contaminants, including excess nutrients [16], microbes [17], sediments [10], and a wide array of anthropogenic chemicals that impact water quality [18]. The latter include contaminants of emerging concern (CECs), a diverse group of synthetic or natural compounds, including pharmaceuticals, personal care products, pesticides, and industrial chemicals [18, 19] that are not commonly monitored or regulated, but are increasingly considered to pose risks to human health and ecosystems [20, 21]. Wastewater represents a significant source of CECs, especially in coastal regions with inadequate waste treatment facilities [22–24]. These include locations that lack treatment altogether (e.g., direct discharge) and those with conventional treatment plants relying on primary (physical settling) and secondary (biological) treatment, which target suspended particles, nutrients, and bulk organic matter but are generally not designed to remove CECs from wastewater [20–22, 25]. Because water quality monitoring programs targeting CECs are relatively uncommon in tropical regions, and toxicological studies using coral reef organisms are rare, our understanding of their ecotoxicological impacts in these environments is still largely lacking [26].

Three interconnected challenges concerning coral reef ecotoxicology underlie this knowledge gap. First, how chemical pollutants disperse within interconnected coastal waters, and which ecotoxicological effects follow, remains poorly understood. Land-derived pollutants are transported, transformed, and delivered to nearshore ecosystems along the ridge-to-reef continuum [27]. Here, they can deteriorate water quality and adversely affect local ecosystems, such as mangroves and seagrass beds, as well as the adjacent fringing coral reefs [10, 28]. The delivery of these pollutants to reefs is modulated by rainfall and local oceanographic conditions [29], and coastal features that restrict circulation, such as semi-enclosed bays, can further shape it by retaining pollutants and releasing them episodically [30]. Whether such variation in pollution produces discrete pulses of elevated exposure on reefs and which ecotoxicological effects this has on reef biota remain largely unknown.

Second, there is a limitation in estimating the realistic environmental exposure concentrations of CECs in coral reef waters, as they often occur in low ambient concentrations (ng L^−1^ to µg L^−1^) and are episodically released in coastal systems [31, 32]. Such episodic events are rarely captured by conventional water quality monitoring, which depends on discrete grab samples collected weeks to months apart [31, 33]. Discrete water sampling may lead to a significant misestimation of both the short- and long-term exposure of marine life to CECs [31]. To address this issue, time-integrated passive sampling devices that accumulate passing contaminants from the water column over extended deployment periods (weeks to months), like the exposure experienced by a sessile organism, could represent a potential solution [31, 34, 35].

Third, water quality assessment frameworks for both freshwater and marine systems are largely based on toxicity data for single substances, which do not capture the combined effect of the many contaminants that co-occur in the environment [26]. Chemicals are commonly present in water as complex mixtures, which can have additive, synergistic, or antagonistic effects, meaning that their combined toxicity can deviate substantially from the sum of that of individual compounds [26]. Chemical-by-chemical risk assessments that compare measured environmental concentrations against ecotoxicological thresholds for single substances are therefore likely to misestimate the cumulative ecotoxicological risks posed by complex environmental pollutant mixtures [36–38]. Estimates of the combined toxic pressure of all bioavailable chemicals present in coral reef environments affecting local biota can be achieved either by computational mixture toxic pressure calculations based on measured chemical concentrations or by direct measurements of the toxic potency of the mixtures using effect-based methods (EBMs). These include a variety of *in vivo* (whole organism) and *in vitro* (cell-based) bioassays to measure biological responses to environmental samples, representing a variety of toxicological modes of action, i.e., physiological or behavioral signs characterizing adverse biological responses [39–41]. Both approaches can be applied to the same passive sampler extracts, enabling a holistic and time-integrated assessment of chemical exposures and the associated ecotoxicological risks [31].

The tropical Southern Caribbean island of Curaçao presents a fitting setting to address this knowledge gap and to overcome the abovementioned interconnected challenges. Recent work has shown that the intensity of human activity on the island, expressed as the proportion of human modified land (“altered surface”, i.e. roads, buildings, and agriculture) in coastal watersheds, correlates with increased input of land-derived pollution to its fringing coral reefs, most strongly at shallow depths and during the wet season [42, 43]. In addition, the many inland bays along the coast of Curaçao have been discussed as potential point sources of water-borne pollutants to the adjacent reefs [30, 44, 45]. However, the identity of the individual CECs (e.g., pesticides, pharmaceuticals, industrial chemicals) entering these bays and reefs has not been determined, nor were their mixture toxic effects in waters along the bay-to-reef gradient.

Therefore, this study aimed to characterize the occurrence and dispersal of anthropogenic chemicals and their potential ecotoxicological effects at three sites with a low, medium, and high proportion of altered coastal surface in their coastal watersheds. Each site comprised an inland bay and its adjacent up- and downstream fringing reefs. Time-integrative passive sampling of dissolved organic chemicals along a gradient from the inland bay to the adjacent up- and downstream fringing coral reefs was conducted during the wet and dry seasons. Chemical mixtures were characterized using high-resolution chemical target analysis, and their toxic potencies were estimated using a combination of compound-based computational chemical risk assessments and a panel of *in vitro* bioassays representing multiple modes of toxic action.

## 2. Methods

### 2.1 Study design and sampling locations

The island of Curaçao has 156,000 inhabitants [46], with ∼1.5 million tourists visiting per year [47] and a surface area of 444 km^2^. It is characterized by a semi-arid climate with a wetter period (October-December) and a drier period (April-June) [48]. Pollutant levels in three inland bays and adjacent reefs were investigated along the island’s leeward coast, with a dominant along-shore northwestern surface current [45] (Fig. 1): Piscadera Bay (0.69 km^2^), Santa Martha Bay (0.73 km^2^), and Awa Blanku (0.10 km^2^).

**Figure 1.**
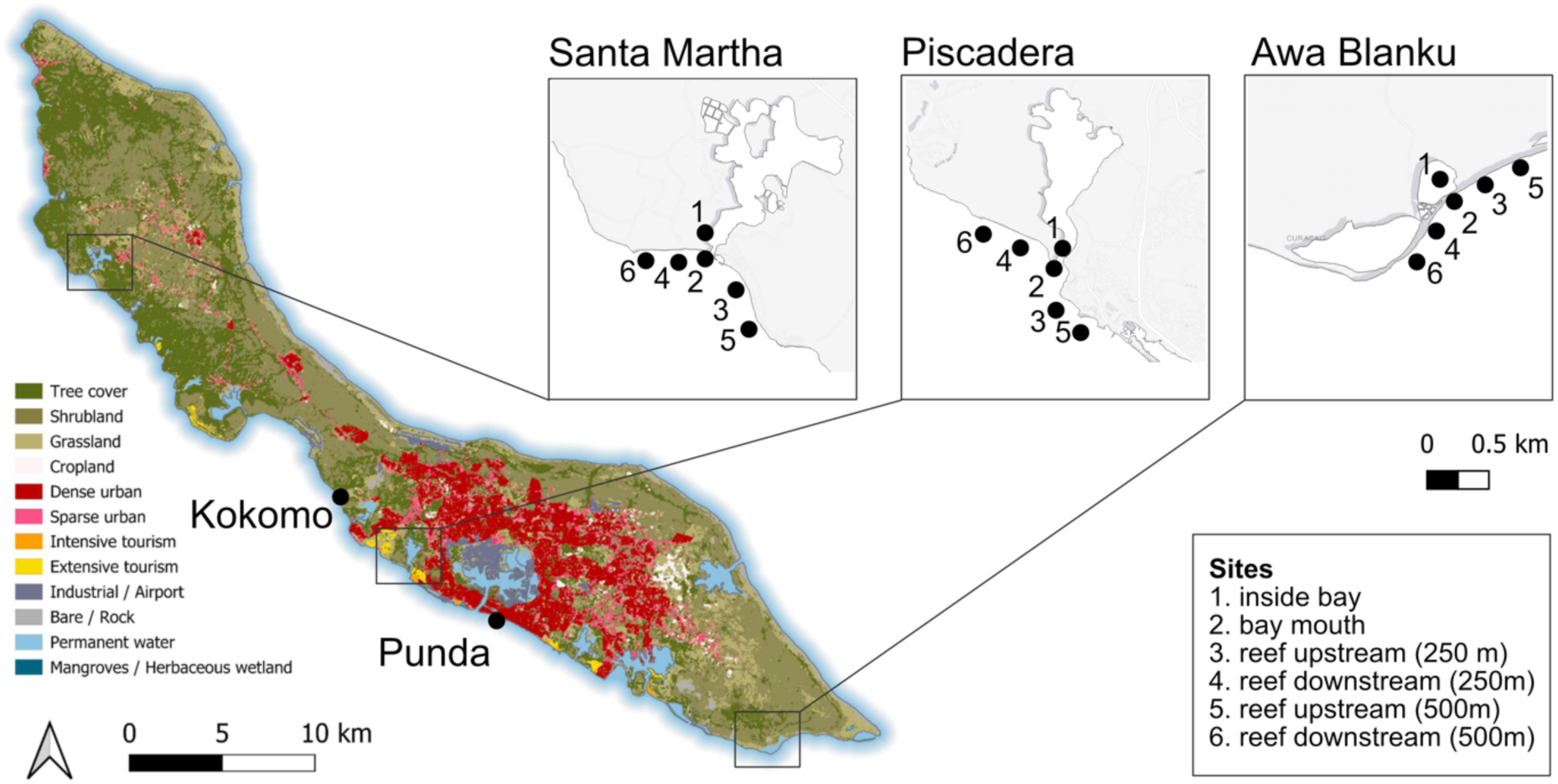
Sampling sites at three inland bays and two additional sites along the leeward coast of Curaçao. Land-use base map from Steward et al. [49]. Sampling campaigns were conducted during the dry season (April-May) and the wet season (October-November) of 2024.

These bays were selected because the proportion of altered surface in their watersheds, and hence the expected intensity of land-based pollution, differs from undeveloped to highly urbanized [42, 43]. Piscadera Bay lies at the western side of Willemstad, the capital city of the island, and serves as an important catchment area for a 13.5 km^2^ watershed in the highly-developed center of the island [44]. A relatively small opening (76-105 m wide; < 6 m depth) connects the bay to nearby reefs. Piscadera Bay is hypothesized as the “highly” polluted site. Santa Martha Bay is considered the “medium” polluted site, as the bay lies towards the less densely populated northwestern side of the island with a catchment area of 16.8 km^2^, and is characterized by a narrow and shallow mouth (40 m wide; < 4 m depth). Awa Blanku Bay is located near the southeastern tip of the island, which was never developed, and is therefore hypothesized as the “low” polluted site [42, 43]. This bay’s mouth is approximately 44 m wide and < 2 m deep, and it is a catchment for 5.7 km^2^ of land. Sampling occurred in the dry season from April 26^th^ to June 1^st^, 2024, and in the wet season that same year from October 31^st^ to December 10^th^.

Along a gradient from bay to reef, six sites were sampled for each bay: inside the bay (1), in the bay mouth (2), and four sites on adjacent reefs up and down the coast at approximately 250 (upstream 3; downstream 4) and 500 m away from the mouth (upstream 5; downstream 6; Fig. 1). The sample depths inside the bay and bay mouth depended on the topography of each bay, between 5–11 m water depth. Water sampling at the reefs was always conducted at 9–11 m water depth. Two additional reef sites, Kokomo and Punda, located away from the bays but also hypothesized as highly polluted, were also included in the study. Punda comprises the eastern coastal section of Willemstad, characterized by the most densely developed urban zones on the island [42, 43]. Kokomo is located downstream from the city and is expected to be exposed to high land-derived pollution as its watershed harbors the island’s landfill [42].

### 2.2 Time-integrative sampling of organic chemicals from the water

#### 2.2.1 Sampling design

To obtain time-integrated samples of organic chemical mixtures in the water column, polar organic chemical integrative passive samplers (POCIS) were deployed at each sampling site. The amount of a compound accumulated in a passive sampler depends on the sampler design (sorbent type and mass, membrane, and exposed surface area), the deployment time, compound properties (e.g. polarity and molecular size), and the local environmental conditions (e.g. water flow, temperature and biofouling of the membranes) [49–51]. Furthermore, properties of the sampler determine which compounds are captured: the membrane (often made of polyethersulfone [PES]) slows the uptake of hydrophobic compounds (log Kow > 2) [49, 51, 52], and different sorbents retain a partly different suite of bioactive compounds, so the choice of sorbent also determines which chemicals and their resulting toxic effects are captured [49]. In addition, the pollutant load and composition of the water matrix (e.g. dissolved organic matter, salinity) can affect uptake by occupying part of the sorbent’s binding capacity or altering the membrane boundary layer which is an effect of the deployment environment rather than of the sorbent itself [51, 53, 54]. At high pollutant loads and extended exposure durations, the binding capacity of the sorbent (saturation) can be approached, after which uptake is no longer linear in time [51]. However, with the POCIS configuration applied here (see section 2.2.2) and month-long deployments, uptake of polar compounds is expected to remain linear [55] and saturation is unlikely at the concentrations found at the study sites, though this was not verified. Hydrodynamic conditions can differ strongly between locations, but this is only rarely considered in passive sampling studies. To overcome this, passive flow monitors (PFMs; see section 2.2.3) were used to account for differences in water movement among sites. Moreover, extract enrichment was corrected for sorbent loss during processing by weighing the remaining sorbent after extraction. This way, effects of site-specific conditions and sampler handling on the study outcomes were accounted for. These corrections were combined in the relative enrichment factor (REF) calculation described in section 2.2.6.

Estimating target compound water concentrations with the use of passive sampling requires correction for compound-specific sampling rates (Rs), as uptake kinetics differ between compounds [40, 51, 56]. This provides time-weighted average target compound concentrations in the water throughout the exposure of the passive sampler. Conversely, for effect-based measurements on passive sampler extracts, the extracts are assessed as an environmental mixture that includes compounds not identified by target analysis, for which no compound-specific Rs can be assigned. For these, a single average Rs for the whole mixture is more appropriate to express the bioassay responses related to surface water levels [50]. Hence, the most appropriate passive sampler extract enrichment correction for each analytical method was applied here to provide an as accurate as possible exposure scenario at the investigated sites.

#### 2.2.2 Passive sampler construction

The passive sampler was designed and constructed based on established methodology [49, 51, 55]. The POCIS were constructed at the University of Amsterdam using 200 mg of Oasis hydrophilic-lipophilic balance sorbent (HLB; Waters, The Netherlands) from solid-phase extraction (SPE) cartridges. All solvents used in the POCIS assembly were of liquid chromatography (LC) grade (Biosolve, Netherlands). Prior to POCIS assembly, the HLB sorbent was conditioned by sequentially eluting 5 mL of acetonitrile (ACN), methanol (MeOH), and Milli-Q water, and dried under vacuum. The HLB sorbent was enclosed between two PES diffusion-limiting membrane filters (0.1 µm pore size, 90 mm diameter; Sterlitech, USA). PES membranes were washed in a 1:1 MeOH:Milli-Q solution (1 min) and rinsed in pure Milli-Q before POCIS assembly. The membranes and sorbent were sandwiched between two stainless steel rings with an inner opening (54 mm diameter) that were secured with stainless steel nuts and bolts inside perforated stainless steel protective covers to prevent damage to the membranes during deployment. All metal parts of the POCIS were washed with acetone and MeOH before assembly to remove contamination. After assembly, POCIS were stored at 4°C in food-grade Mylar zip-lock bags until transport to Curaçao. Blank POCIS were transported to Curaçao, kept at -20°C during deployment of the other samplers, and processed alongside the exposed samplers to ensure POCIS had remained free of potential pollutants during fabrication and handling before deployment (see section 2.3.1).

#### 2.2.3 Passive flow monitoring for correction to local hydrodynamics

PFMs were prepared based on the method of O’Brien et al. [57]. These simple devices served to roughly estimate the amount of passing water each POCIS had been exposed to during deployment. PFMs were constructed by mixing 300 g dental plaster (Whip Mix Corporation, Louisville, USA) with 160 mL of Type I ultrapure water (Elga Purelab Classic UV). The resulting mix was cast into individual 50 mL plastic specimen containers (dimensions: 40×60x21 mm) and left to dry to constant weight at room temperature. Once dry, the containers were weighed (0.1 g resolution), closed with a screwcap, and stored at room temperature until deployment alongside each POCIS. The relative weight loss of the plaster before and after installation was used as a relative measure for flow at the respective site (see section 2.2.6).

#### 2.2.4 Field deployment of POCIS and PFM

POCIS (*n* = 4) and PFMs (*n* = 3) were deployed on SCUBA at all six sampling locations for each bay and at the two reef sites, Punda and Kokomo (in total, 80 POCIS and 60 PFMs were deployed). Four pieces of rebar (75 cm) were placed approximately 1–3 m apart and driven into the seafloor using a hammer. Each POCIS was secured to one of the rebars using cable ties at approximately 0.5 m above the reef. PFMs were attached next to three POCIS using cable ties. The POCIS and PFMs were deployed for 29–34 days (Table S1), after which they were collected and individually placed in zip-lock bags and transported in the dark to the laboratory on ice. In the lab, they were stored at 4°C until processing. Within 24 h of retrieval, the POCIS were disassembled, and the HLB sorbent was transferred from the membranes into a collection vessel using ultrapure water (15 mL). For samples collected during the dry season, the HLB sorbent of each POCIS was collected in a 15 mL plastic centrifuge tube (VWR, Netherlands). For samples collected during the wet season, it was directly transferred into a solvent-rinsed (ACN, MeOH) SPE column using a glass funnel, after which the columns were sealed with a Luer tip and cap and secured with Parafilm. The sorbent was stored at -20 °C and transported on dry ice to the University of Amsterdam for further analyses.

#### 2.2.5 POCIS extraction

All equipment used in the POCIS extraction was first cleaned with acetone and LC-grade methanol (Biosolve, The Netherlands). For dry and wet season samples, the recovered HLB sorbents were respectively thawed and transferred to solvent-rinsed SPE columns using a glass funnel, or the SPE columns already containing the HLB sorbent were thawed. All SPE columns were placed on an SPE manifold and first eluted with 5 mL LC-grade ACN, followed by 5 mL MeOH (both under gentle vacuum at slow dropwise flow rate), resulting in a 10 mL total extraction volume. All extracts were weighed to correct for any solvent losses during extraction, expressed as the extract recovery. Extracts were stored in 10 mL glass vials at - 20 °C. After extraction, the sorbents were dried and weighed to determine the sorbent recovery relative to the initial 200 mg weight per POCIS to correct for potential loss of sorbent during sampling and processing.

#### 2.2.6 Determination of extract enrichment

The REF was calculated for each POCIS extract to estimate the extract enrichment compared to the surface water. A REF value approximates the sampled water volume after correcting for sorbent recovery, extraction recovery, and site-specific flow conditions. The PFM mass loss was used to account for variation in the amount of water that had passed the POCIS during deployment at each site. The mass loss of each PFM was expressed relative to the average loss across all sites, converting the absolute mass loss into a dimensionless correction factor. This deviation from the average loss was used to adjust the REF, thereby correcting the results for differences in local current velocity.

The REF for each extract was hence calculated as:

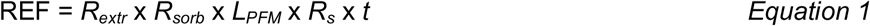

Where *R_extr_* is the extract-recovery, *R_sorb_*is the sorbent-recovery, *L_PFM_* is the relative PFM loss, *R_s_*is the sampling rate per day, and *t* is the exposure time (days of deployment). The *R_s_* used depended on the subsequently applied analytical method. For the target chemical analysis (section 2.3.1), previously reported compound-specific *R_s_* values (Table S2) were used, providing a compound-specific REF for each detected chemical. For the bioassays (section 2.3.2), a single *R_s_* of 0.1 L d^−1^ was used for the whole mixture, because the extracts contain the full chemical mixture, including compounds not identified by target analysis, and bioassay responses are thus best represented by a single sampled volume per sampler. This value was previously applied to bioassay extracts of POCIS deployed in Santa Martha Bay [30] and is representative of the compound-specific *Rs* values reported for polar organic compounds in POCIS [51]. In both cases, the PFM mass loss in the REF corrects the estimated sampled volume for site-specific flow conditions.

To ensure that all replicate extracts from the same site fell within a comparable concentration range for the subsequent analysis, all replicates were diluted to the same REF. For each site, the replicate with the lowest REF was identified, and all other replicates from that site were diluted with 1:1 ACN:MeOH to match that lowest REF. This minimized the risk of poor detectability due to different concentration ranges or matrix effects.

### 2.3 Chemical- and bioanalytical characterization of passive sampler extracts

#### 2.3.1 Target chemical analysis by liquid chromatography-mass spectrometry

POCIS extracts were analyzed for the concentrations of 76 target organic chemicals spanning pharmaceuticals, industrial chemicals, and pesticides (Table S2) based on the method developed by Specker et al. [58]. Of each extract, an aliquot was diluted in Milli-Q and spiked with labelled internal standards to a final concentration of 0.2 µg L^−1^ (Table S3). The blank POCIS extracts and method blanks were prepared and analyzed in the same way. All extracts (20LJμL injection volume) were analyzed using ultra-high-performance liquid chromatography (UHPLC, Nexera 40 series, Shimadzu, Den Bosch, the Netherlands) using a Kinetex® UPLC Biphenyl (1.7LJµm particle size, 100LJÅ pore size, 50LJ×LJ2.1LJmm) chromatographic column coupled to a trapped ion mobility separation (TIMS) quadrupole time-of-flight instrument (timsTOF Pro 2, Bruker Daltonics, Leiderdorp, the Netherlands). A vacuum-insulated probe heated electrospray ionization (VIP-HESI) source was used in both negative and positive ionization modes, operating in the mass range 20–1300LJm/z. Additional details, including liquid chromatography-mass spectrometry (LC-MS) settings, are provided by Specker et al. [58].

The LC-MS data were analyzed using TASQ (Target Analysis for Screening and Quantitation) from Bruker (Version: 2023.1.5 10496) with a mass tolerance of 3LJmDa, a retention time window of ≤ 0.5 min excluding < 1000 intensities, and the collision cross-section values provided by Specker et al. [58], using an extracted ion chromatogram 1/k0 window of 0.035. Calibration curves for all target analytes were prepared in Milli-Q water over a concentration range from 0.03LJµg L^−1^ toLJ16 µg L^−1^ via serial dilution, with each dilution level containing 0.2LJµg L^−1^ of labelled internal standards (Table S3). Calibration curves had an r^2^ > 0.99 with at least seven calibration points and less than 30% residuals (Table S2). Further details on data processing in TASQ are provided in Specker et al. [58]. Previously reported compound-specific REF were used to calculate extract concentrations (C_extr_) for the detected target chemicals using Equation 1 (Table S2). The surface water concentrations (C_w_) for each chemical based on their extract concentrations were calculated using Equation 2:

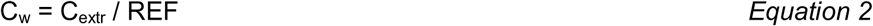

#### 2.3.2 Effect-based mixture toxicity assessment

The effect-based mixture toxicity assessment characterized the ecotoxic potency of the total chemical mixture extracted from the water using passive samplers at a selection of sites during the wet season. These were prioritized for their relatively high seasonal chemical concentrations and value in investigating transport of chemicals from bays to reefs, while restricting the number of analysed extracts to limit the bioanalytical costs. For each bay, this resulted in one sample, constituted from replicate POCIS, from the inside of the bay, one from the bay mouth, and one from an adjacent reef site. The toxic potency of the extracts was assessed by a panel of *in vitro* Chemically Activated LUciferase eXpression (CALUX^®^) bioassays [40, 41]. These assays use engineered human cell lines to detect whether compounds in a mixture activate or block specific receptor pathways [39]. They provide a standardized measurement of bioactivity in the sampled mixture rather than organism-specific toxicity [39, 41]. The CALUX assays were used to quantify agonism of the estrogen receptor (ERα), aryl hydrocarbon receptor (AhR; PAH CALUX, referred to as PAH below), and pregnane X receptor (PXR), as well as antagonism of the androgen receptor (*anti*-AR). Additionally, oxidative stress response was quantified with the nuclear factor erythroid 2-related factor 2 (Nrf2) CALUX assay [59], and cytotoxicity was measured as a general indicator of cell health and to verify that the other bioassay responses were not a result of reduced cell viability [59, 60]. These endpoints together capture major toxicological pathways that are commonly affected by complex pollutant mixtures in water [39, 40]. For the bioassays, all replicate extracts per site were pooled, evaporated to dryness under a stream of nitrogen, and redissolved in dimethyl sulfoxide (DMSO) at the BioDetection Systems laboratory (Amsterdam, The Netherlands), where all CALUX assays were performed at a final concentration of 0.1-1.0% DMSO. Bioassay responses were expressed as bioanalytical equivalent concentrations (BEQ), i.e., the concentration of a reference compound that produces the same response in the assay as the sample [41]. BEQs were converted to surface water values (BEQ_W_) by dividing them by the REF of the pooled extract (section 2.2.6) using Equation 3:

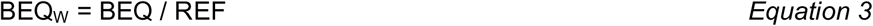

### 2.4 Ecotoxicological risk assessment using multiple lines of evidence

#### 2.4.1 Risk assessment using individual chemical thresholds

An ecotoxicological risk assessment was performed for each quantified target compound at each site and season. Compound concentrations derived from the POCIS were compared to predicted no-effect concentrations (PNECs) for marine water, applicable to the sampled coastal sites, obtained from the NORMAN Ecotoxicology Database [61]. A PNEC is the concentration below which no adverse effects on sensitive aquatic organisms are expected. Risk quotients (RQs) were calculated by dividing the concentration of each target compound by its PNECs. The NORMAN PNECs are derived from experimental acute and chronic toxicity data, or where these are lacking, from quantitative structure-activity relationship (QSAR) predictions using assessment factors following European guidance [62]. Marine PNECs are largely extrapolated from freshwater data and are not specific to tropical waters. Toxicity thresholds derived for tropical marine species exist for very few CECs [26, 63, 64], so the marine PNECs represent the best available benchmark for this assessment. The individual RQs per site were added to obtain a summed risk quotient (ΣRQ) as a screening metric that integrates individual compound risks under the concentration addition assumption [37, 38]. This does not predict actual mixture toxicity or account for synergistic or antagonistic interactions, but it provides a relative ranking of sites by their summed chemical risk.

#### 2.4.2 Mixture toxicity assessment using multi-substance toxic pressure

The multi-substance potentially affected fraction of species (msPAF) [65] was calculated from the target compound concentrations at each site and season using the calculator provided by the Dutch National Institute for Public Health and the Environment (RIVM). The calculation uses species sensitivity distributions (SSDs) [66, 67]. An SSD is a statistical distribution of effect concentrations of a chemical for multiple species (minimum of 5 to 10 distinct species from multiple taxonomic or trophic groups, e.g., algae, invertebrates and fish). The fraction of that distribution that lies below the measured concentration is the potentially affected fraction (PAF) of species for that compound. Single PAFs for multiple chemicals are combined into a single msPAF per site, assuming concentration addition for compounds sharing the same toxic mode of action and response addition across different modes of action [65]. SSDs, which are predominantly based on freshwater species [66, 67], were available for 14 of the 24 quantified compounds, and the msPAF was therefore calculated for this subset. Hence, the following compounds were not included: DEET, desvenlafaxine, fluconazole, lamotrigine, naproxen, sucralose, TCPP and venlafaxine.

#### 2.4.3 Mixture toxicity assessment using in vitro bioassays

The CALUX bioassay responses were used as indicators of the potential of the chemical mixture to exert toxicity via the different modes of action studied [39]. In order to distinguish between acceptable and poor water quality, the bioassay responses were compared to effect-based trigger (EBT) values derived from literature [39]. EBT values are bioassay-specific thresholds: when a sample exceeds the EBT, there is a likelihood of toxic effects (i.e., risk) on aquatic organisms [41]. As the sensitivity of coral reef organisms to chemical toxicity is poorly studied and the available EBTs were developed for temperate freshwater species, a precautionary assessment factor of 10 was applied to lower the EBT values, in line with the extrapolation factors proposed for applying temperate toxicity data to tropical species [68]. The bioassay responses were divided by their corresponding precautionary EBT values to calculate effect-based risk quotients (EBRQ; [40]) and summed per site to the cumulative ecotoxicological risk (CER) [31].

### 2.5 Data analyses

All statistics were performed in R [69] using RStudio [70], in which multivariate analyses were carried out in the “vegan” package [71]. Differences in chemical mixture composition between bays, sites and seasons were tested with permutational multivariate analysis of variance (PERMANOVA; [72]), which tests whether the centroids of the groups differ in multivariate space using permutations of the observations. Permutational analysis of multivariate dispersions (PERMDISP; [73]) tests whether the spread of observations around those centroids differs between groups. PERMDISP was run alongside PERMANOVA because PERMANOVA is sensitive to unequal dispersion: a significant PERMANOVA could reflect differences in within-group variability rather than in composition. Principal component analysis (PCA) was used to visualize the patterns tested, and redundancy analysis (RDA) to relate bioassay responses to compound concentrations.

The analyses were based on the 24 compounds for which compound-specific POCIS *R_s_* were available and that were therefore quantified (of the 76 target compounds, 28 were detected; section 3.1). Each of the four replicate POCIS per site and season was treated as one observation, so that the site-level tests are based on the variation among replicates. Concentrations were log-transformed [log10(x + 1)] to reduce the influence of extreme values and non-detects and scaled to unit variance so that the analyses reflect mixture composition rather than compounds with the highest concentrations. Pairwise Euclidean distances were computed from this matrix and used for both PERMANOVA (adonis2, 9999 permutations) and PERMDISP (betadisper followed by permutest, 9999 permutations).

For the season by site interaction, permutations were restricted to within site, because sites were sampled repeatedly across seasons. The following models were tested: (a) the season by bay interaction within the three bays; (b) the season by site interaction; (c) bay differences within each season (controlling for site); (d) inside bay versus adjacent reef composition within each bay and season; (e) differences among reef sites across the island, including the two reef sites without adjacent bays (Punda and Kokomo), within each season. For models (d) and (e), pairwise PERMANOVAs were run and because each set of pairwise comparisons inflates the chance of false positives, the p-values were corrected for multiple testing using the Benjamini-Hochberg false discovery rate (FDR) method [74]. Effects were considered significant at p < 0.05.

Where dispersion differed significantly between groups (PERMDISP), PERMANOVA effects were interpreted with caution, because differences in mixture composition and in within-group variability cannot be fully separated.

PCA was used to visualize mixture composition among sites and seasons, using the same log-transformed data as used for PERMANOVA. Replicates were averaged to station centroids (Fig. 3). Redundancy analysis (RDA) was performed to examine the relationship between the compound concentrations (predictors) and the bioassay responses (response variables) of the wet-season sites for which bioassay data were available.

## 3 Results and Discussion

### 3.1. Occurrence and composition of target compounds

Of the 76 target compounds (Table S2), 28 were detected at least once in the POCIS extracts, while 48 were below the limit of detection (Figs. 2 and S1; Tables S2 and S5). Of the 28 detected compounds, only 24 were quantified. This is due to the absence of compound-specific POCIS sampling rates for fenuron, neburon, pyridafol, and salicylic acid.

**Figure 2.**
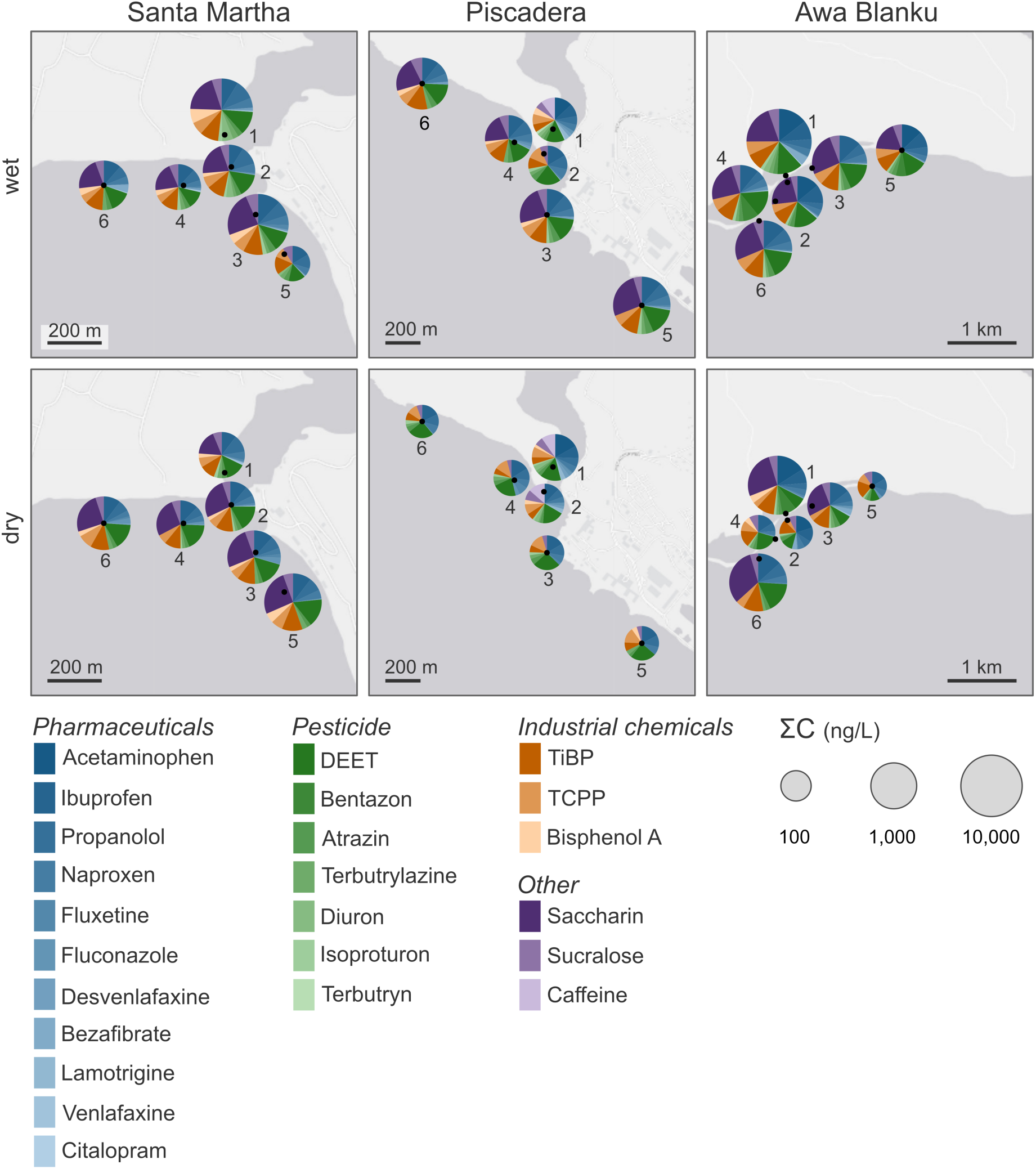
Spatial and seasonal concentrations of target compounds from passive sampler extracts of inland bays and adjacent coral reefs on Curaçao, southern Caribbean. For each bay, pie charts show the 24 quantified compounds collected by polar organic chemical integrative samplers (POCIS) and detected at each site during the wet (top) and dry (bottom) seasons of 2024. Pie size represents the total summed concentration of all compounds (ΣC, ng L^−1^, log scale), and slices show individual compounds (log_10_ (1+C)), colored by compound group.

**Figure 3.**
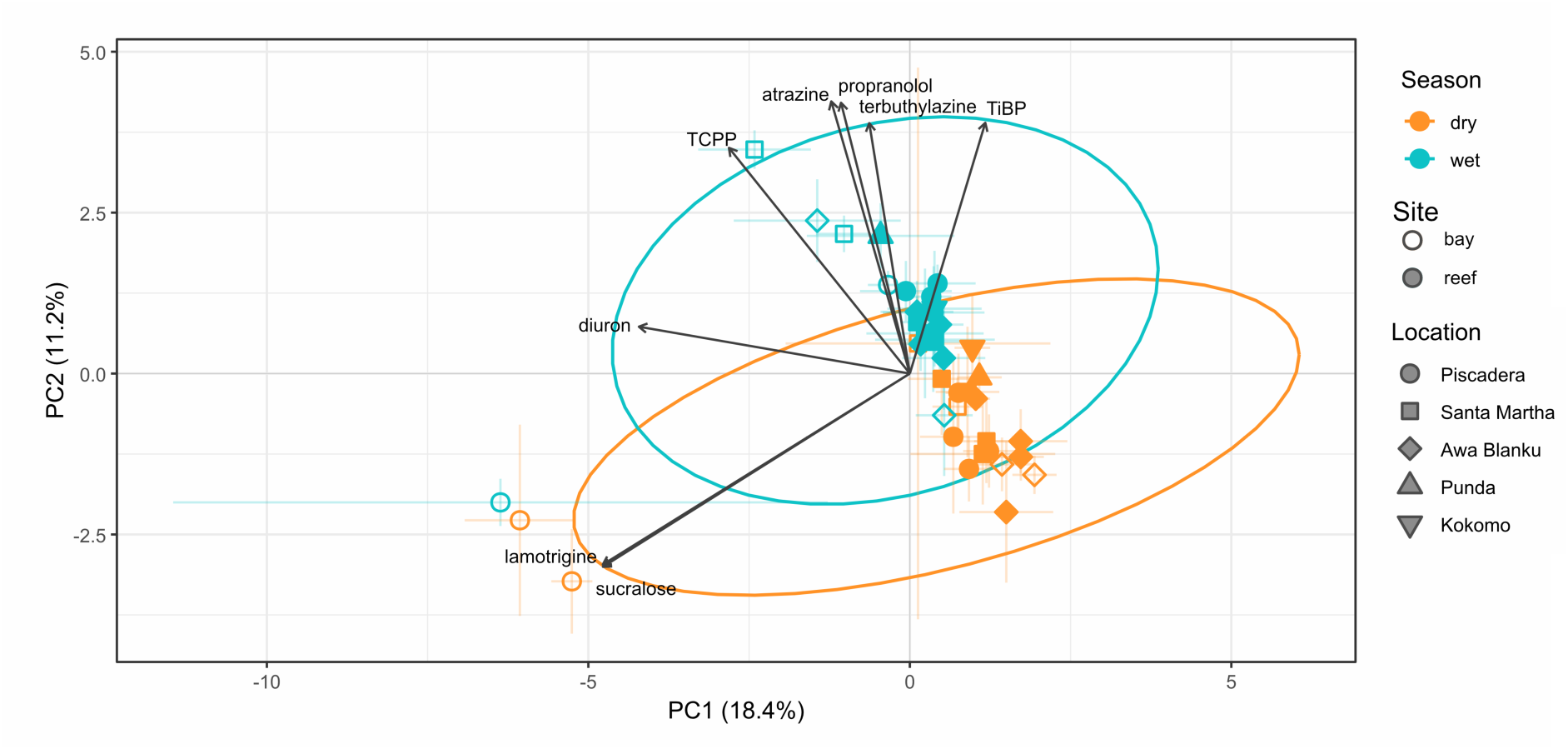
Principal component analysis (PCA) of chemical concentration profiles across sampling sites surrounding three inland bays in Curaçao during two seasons (wet and dry). Colors represent season (orange = dry, blue = wet) and shapes represent site groups. Ellipses indicate 95% confidence regions for each season. Arrows show compound loading vectors with a combined loading of > 0.2. Compound groups (of the 24): Pharmaceuticals (11 compounds; lamotrigine, propranolol), Pesticides (7; diuron, atrazine, terbuthylazine), Industrial chemical (3; TCPP, TiBP), and Other (3; sucralose).

#### 3.1.1. Concentrations peak inside bays in the wet season

Cumulative concentrations were highest inside the bays during the wet season, in every bay (Table 1; Table S6; Fig. 2). Awa Blanku Bay reached a total detected chemical concentration of 1111 ng L^−1^, more than twice that of any reef value, of which nearly half (44%) was the single compound DEET (insect repellent; section 3.1.3). In the dry season, the ranking of the bays reversed and followed the gradient in coastal development: Piscadera Bay had the highest cumulative concentration, followed by Santa Martha Bay, and Awa Blanku Bay (Table 1). Awa Blanku Bay thus showed the largest seasonal difference (971 ng L^−1^), whereas Piscadera Bay was more stable (76 ng L^−1^ difference between seasons; Table 1). In the wet season, the mixtures inside all three bays were dominated by pharmaceuticals and pesticides (45-59% of the cumulative concentration; Table 1), with DEET as the main pesticide. Averaged over the three bays, pesticide and pharmaceutical concentrations inside the bays were 3.0 and 2.4 times higher, respectively, than those at the adjacent reef sites (Table S6) with the largest bay-to-reef differences found for pesticides at Santa Martha Bay (5.7x).

**Table 1.** Cumulative water concentrations (ng L^−1^; sum of 24 quantified compounds) per site along the bay-to-reef gradient and season. Reef values are the mean (range) of the four reef sites per bay. Punda and Kokomo are reef sites without an adjacent bay sampled. The difference between wet and dry seasons and the ratio refer to the site inside the bay. The dominant compound group inside the bay is given with its percentage of the total cumulative concentration.

| Bay | Season | Inside bay | Bay mouth | Reef mean (range) | Wet – dry inside bay (ratio) | Dominant group inside bay (% of cumulative conc.) |
|---|---|---|---|---|---|---|
| Santa Martha | wet | 734 | 384 | 280 (227-348) | 404 (x2.2) | pesticides (45%) |
|  | dry | 329 | 179 | 233 (171-305) |  | pesticides (41%) |
| Piscadera | wet | 464 | 331 | 320 (285-421) | 76 (x1.2) | pesticides (59%) |
|  | dry | 388 | 302 | 158 (130-190) |  | pesticides (60%) |
| Awa Blanku | wet | 1111 | 275 | 370 (325-446) | 971 (x7.9) | pesticides (45%) |
|  | dry | 140 | 93 | 153 (115-192) |  | industrial (43%) |
| Punda | wet |  |  | 475 |  |  |
|  | dry |  |  | 236 |  |  |
| Kokomo | wet |  |  | 436 |  |  |
|  | dry |  |  | 319 |  |  |

#### 3.1.2. Each bay has a unique chemical mixture

The three bays differed significantly in mixture composition, and this difference depended on season (PERMANOVA; Table S4). Within each season, the three bays remained distinct, but more strongly in the dry season (R^2^ = 0.17) than in the wet season (R^2^ = 0.09), and the seasonal shift in composition itself varied among sites within each bay (Table S4).

Composition also separated the inside-bay sites from their adjacent reefs, in every bay and season (PERMANOVA, all *p_adj_* < 0.03; Table S4), but most strongly in Piscadera Bay. The reefs themselves were not uniform: mixture composition differed among all reef sites in both seasons, including the two reefs without an adjacent bay, Punda and Kokomo (reef-only PERMANOVA; Table S4).

Mixtures inside the bays were more heterogeneous than those at the adjacent reefs (PERMDISP; Table S4). Because PERMANOVA is sensitive to such differences within-group dispersion, part of the bay-reef separation described above may reflect the greater variability among bay samples rather than a shift in average mixture composition alone (section 2.5). The greater variability within bays is consistent with bays receiving more variable land-based pollution inputs (e.g., wastewater pulses, seasonal runoff), leading to a chemically heterogeneous mixture [10, 30, 44]. In contrast, reef sites are more exposed to mixing with open ocean water, which is expected to dampen local variability [44, 45, 75].

#### 3.1.3. Individual targeted chemical compounds

The PCA of the 24 quantified compounds visualizes these spatial and seasonal patterns (Fig. 3). Its first two axes explained 18.4% and 11.2% of the total variance, respectively (Table S7). Along PC1, the compounds lamotrigine, sucralose, and diuron pointed in the same direction, separating the inside of Piscadera Bay from the other sampling sites. PC2 contrasted two mixture types: pesticides (atrazine and terbuthylazine) and organophosphate flame retardants (TCPP and TiBP) together with propranolol at one end, and the wastewater tracers sucralose, lamotrigine and fluconazole at the other. Wet-season samples were located towards the pesticide and industrial chemical end of PC2, separated from the dry-season samples, consistent with the season-by-bay interaction (Table S4). The Piscadera inside-bay samples separated most clearly along the wastewater-tracer axis.

The sites were distinguishable by two groups of chemicals (Fig. 3; Table S7). Piscadera Bay was distinguished by the persistent wastewater tracers sucralose, lamotrigine, and fluconazole in both seasons, indicating a continuous domestic wastewater input in that bay. The wet-season samples of all bays were enriched in pesticides and organophosphate flame retardants (atrazine, terbuthylazine, triisobutyl phosphate) together with propranolol. This is consistent with increased surface runoff [10] and with sewer overflows, as Curaçao’s combined sewer system discharges sewage and rainwater into the ocean during heavy rainfall [42, 76, 77]. During the wet season, increased outflow from the bays transports bay-derived pollutants towards the reefs, where they undergo dilution and mixing [44]. In the dry season, the water exchange between the bay and the ocean is more limited [44], resulting in more uniform chemical mixtures less influenced by local inputs into the coastal waters in which the reefs are located.

The wastewater signature was reinforced by the artificial sweeteners. Sucralose, which is highly resistant to degradation and is a well-established tracer for WWTP effluent [23], was detected in Piscadera Bay. Contrastingly, saccharin, which is a more labile sweetener that is typically removed with high efficiency (>90%) during wastewater treatment and degrades in the marine environment [78], was not detected (limit of detection: 7.55 ng L^−1^). The presence of persistent sucralose alongside non-detection of the more labile saccharin is consistent with a sustained treated wastewater signature in Piscadera Bay.

A seasonal runoff signal was exemplified by the insect repellent DEET, which reached the highest concentrations of any quantified compound and strongly influenced differences in cumulative concentration (section 3.1.1). Its concentration varied most in Awa Blanku Bay, from 530 ng L^−1^ in the wet season to 8.6 ng L^−1^ in the dry season (Table S5), characteristic of increased repellent use and runoff in the wet season. The concentrations measured in the wet season were substantially higher than those reported in other Caribbean studies (e.g., up to 49 ng L^−1^ in the Florida Keys [79]).

### 3.2. Ecotoxicological risk assessment across sites and seasons

#### 3.2.1. Risk assessment using individual and summed chemical thresholds

RQs translate the measured concentrations into potential ecotoxicological effects. Of the 24 quantified compounds, 22 exceeded an RQ of 1 in at least one sample, meaning that their measured concentration surpassed the PNEC and effects on sensitive organisms cannot be excluded at these sites (Fig. 4). Individual RQs and their sum (∑RQ) were highest inside the bays during the wet season, pointing to proximity to land-based sources, limited dilution in the inland bays, and rainfall-driven input as the main drivers of risk for the target compounds. The herbicide diuron, which reaches water through run-off after application to crops [80], had substantially higher RQs during the wet season inside all three bays and Punda (Fig. 4).

**Figure 4.**
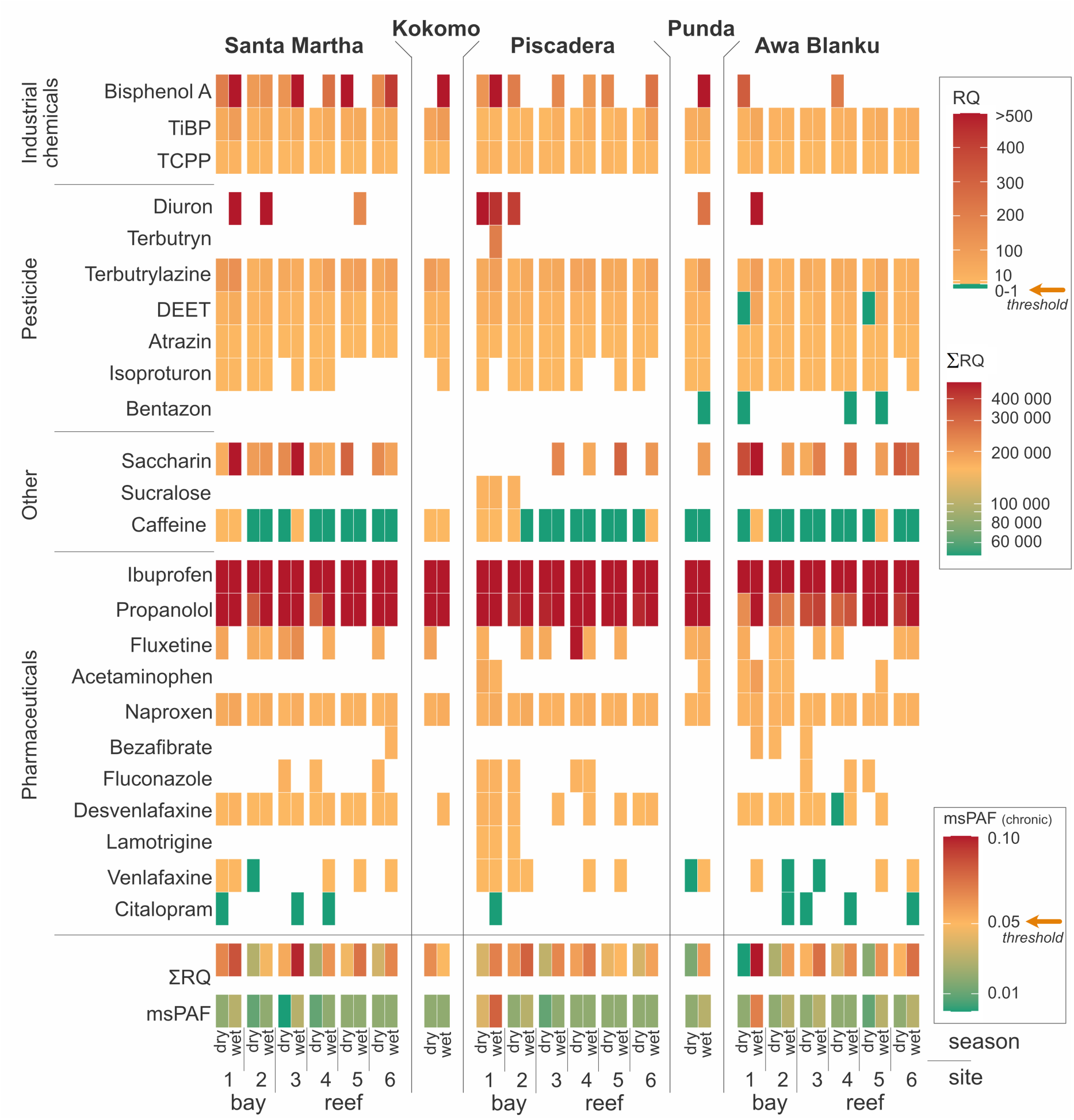
Heatmap of ecotoxicological risks across bays and seasons for target chemicals. Compound-specific risk quotients (RQ), calculated using predicted no-effect concentrations (PNECs) for each site in the dry and wet season for the three bays in Curaçao (Piscadera, Santa Martha, and Awa Blanku). Compounds are grouped by class (industrial chemicals, pesticides, other, pharmaceuticals). RQ values <1 are shown in green, and yellow to red represent values >1 and <500. The green to red color spectrum indicates the ∑RQ values between >60 000 and <400 000. The bottom row shows the chronic multi-substance potentially affected fraction of species (msPAF chronic) per site and season, with a green to red color spectrum indicating the values between 0.00 and 0.10.

A subset of compounds was consistently associated with the highest risks, notably the pharmaceuticals propranolol and ibuprofen (Fig. 4). The beta-blocker propranolol can alter heart rates, behavior, and reproductive functions in fish and invertebrates [81, 82]. Similarly, ibuprofen can induce oxidative stress and disrupt developmental and reproductive functions in aquatic organisms [83, 84]. While specific data on reef organisms are scarce, the presence of these biologically active compounds raises concern, as tropical reef organisms may be more sensitive to chemical pollutants compared to temperate marine or freshwater organisms [68, 85].

In contrast, other compounds like the herbicide bentazon and the antidepressant citalopram were consistently detected at RQs below 1, suggesting minimal direct risk to reef biota. Bentazon is a herbicide primarily used for soybeans, rice, corn, mint, and peanuts [86], none of which are cultivated on Curaçao. However, bentazon is also applied for weed control in turf and ornamental planting, and it is highly water-soluble and mobile [86], so non-agricultural use in gardens, resorts, or golf courses is a plausible local source, and its consistent detection at low concentrations is more in line with diffuse, low-intensity inputs.

#### 3.2.2. Mixture toxicity assessment using multi-substance toxic pressure

The msPAF confirmed that the highest cumulative toxic pressure occurred inside the bays during the wet season (Fig. 4). The highest chronic msPAF values were observed during the wet season inside Piscadera Bay (8%) and Awa Blanku Bay (7%), both exceeding the 5% threshold for the affected fraction of species, commonly used to indicate a level of concern for ecosystem protection [66]. In Santa Martha Bay, the chronic msPAF was lower, remaining below this threshold (3% wet, 2% dry season). On the reefs, msPAF values ranged from 1–3%, indicating consistently lower toxic pressure than inside the bays. This decreasing bay-to-reef pattern was most evident at Piscadera Bay, where during the wet season, the chronic msPAF decreased from 8% inside the bay to 3% at the mouth and 1–2% on the adjacent reefs. However, these values are conservative, as SSDs were not available for all compounds (14 out of 24; section 2.4.2), and frequently detected compounds (DEET, desvenlafaxine, fluconazole, lamotrigine, naproxen, sucralose, TCPP and venlafaxine) could not be incorporated into the msPAF calculation, despite their presence.

#### 3.2.3. Strongest bioassay response on reefs

Bioassays were performed on the pooled wet-season extracts of the inside-bay site, the bay mouth, and one reef site of each bay (section 2.3.2), from which several patterns emerged (Fig. 5). First, the oxidative stress response (Nrf2) was by far the strongest and, unlike the target compound concentrations, which decreased from the bays to the reefs, it was the highest at the reef sites. It dominated the CER, which therefore also peaked on the reefs. Second, the anti-AR response was only found at Awa Blanku Bay, inside the bay and at its mouth. Third, ERlll, PAH, and PXR responses were low or absent at all tested sites, and no cytotoxicity was found. The two lines of evidence thus answer different questions and are complementary rather than contradictory: the target compound analysis identifies the bays as sources and reservoirs of the chemicals, whereas the effect-based assessment identifies the reefs as sites where the mixture is most biologically active.

**Figure 5.**
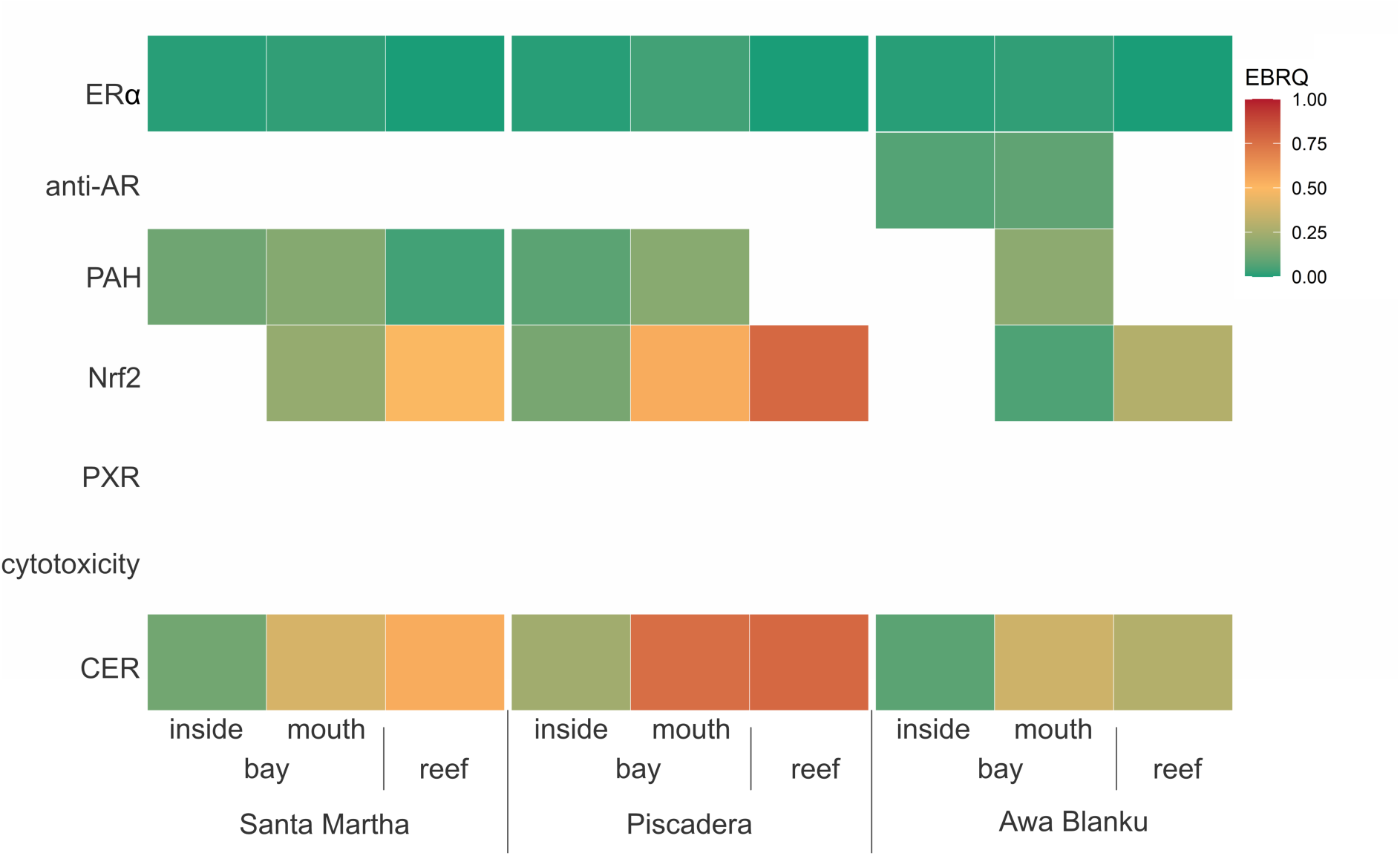
Heatmap of in vitro bioassay responses across bays during the wet season. Effect-based risk quotients (EBRQ) for three sites (inside bay, bay mouth, reef) at three bays in Curaçao (Santa Martha, Piscadera, and Awa Blanku). Estrogen receptor = ERα), androgen receptor antagonism = anti-AR, aryl-hydrocarbon receptor = PAH, oxidative stress response = Nrf2, pregnane X receptor = PXR, and the cumulative ecotoxicological risk = CER. The color scale indicates the level of risk: EBRQ/CER = 0-1.

An RDA relating the bioassay responses to the compound concentrations shows how the two lines of evidence align (Fig. 6). The first RDA axis separates the sites along the Nrf2 gradient, and the Nrf2 response correlates with the herbicide atrazine and the antidepressant fluoxetine. The anti-AR vector points in the opposite direction and correlates with the DEET concentration.

**Figure 6.**
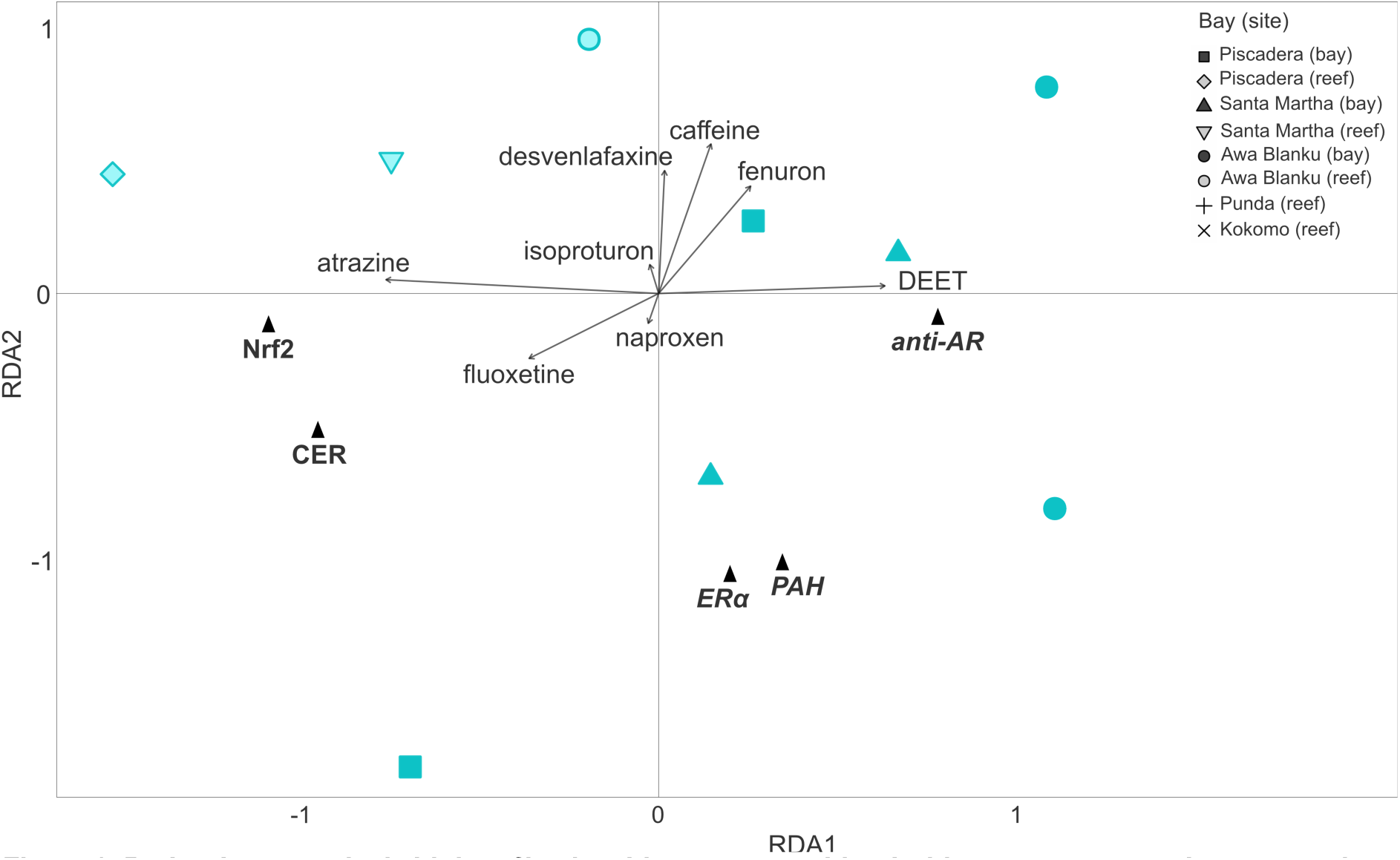
Redundancy analysis biplot of in vitro bioassays combined with target compound concentrations for three sites at three bays in Curaçao (Piscadera (1), Santa Martha (2), and Awa Blanku (3)) during the wet season. The bioassays are depicted as triangles. Estrogen receptor = ERα, androgen receptor antagonism = anti-AR, aryl-hydrocarbon receptor = PAH, oxidative stress response = Nrf2, pregnane X receptor = PXR, and the cumulative ecotoxicological risk = CER. Vectors (arrows) of only significant compounds.

The Nrf2 pathway, which mediates the oxidative stress response, is a cellular defense mechanism that can be activated by various compounds, including metals, pesticides, and other organic micropollutants [87, 88]. The two compounds correlated with the response, atrazine and fluoxetine, are possible contributors, as atrazine can induce oxidative imbalance in fish [89] and fluoxetine can cause oxidative stress in aquatic organisms, from fish [90, 91] to marine macroalgae [92]. Most of the oxidative stress response in water samples is usually caused by chemicals that are not identified by target analysis [60]. Hence, atrazine and fluoxetine are likely proxies for a wider wastewater- and runoff-derived mixture, which was largely overlooked by the applied target chemical analysis, but may well induce the observed Nrf2 response. Reefs experience high levels of recreational activity during which chemicals from sunscreens and other personal care products are released directly into the water column by swimmers and divers [93]. The high penetration of UV radiation in clear reef waters can accelerate photodegradation of pollutants into more reactive transformation products [94], which may partly explain the relatively high observed Nrf2 responses to reef site extracts. Additionally, the reef itself is a source of bioactive compounds: corals, sponges and algae produce a diverse range of secondary metabolites for defense, competition and signaling, including terpenoids, pro-inflammatory oxylipins, and other reactive lipids [95, 96], which may have also contributed to the observed oxidative stress signal. Hence, the relatively high response to the reef site extracts may also be partly attributable to naturally occurring allelochemicals rather than to anthropogenic pollutants alone. Given the lack of cytotoxicity detected in any of the samples, the observed Nrf2 response is not an artifact of general cellular stress induced by the extracts.

Anti-androgenic activity, measured via the anti-AR CALUX, was only detected inside and at the mouth of Awa Blanku Bay. Known androgen receptor antagonists in surface waters include several pesticides and ingredients of personal care products such as UV filters and parabens [40, 97]. The co-occurrence of the anti-AR response with the highest DEET concentrations at Awa Blanku Bay (Fig. 6) points to the use of personal care products as a plausible local source. However, DEET itself is inactive in androgen receptor antagonist assays [98]. Hence, in the present study, DEET likely acts as a tracer of personal care product use, which contain anti-androgenic ingredients such as UV filters and parabens [97]

There was a low to no response for the ERα, PAH, and PXR bioassays to all extracts. ERα activity is a well-established indicator of wastewater-derived estrogens [40, 99] and was low at all sites (EBRQ below 1), in line with De Jong et al. (2025) [30], who found similarly low ERα activity in Santa Martha Bay. The estrogens that commonly drive ERα activity in wastewater (estrone, 17β-estradiol, 17α-ethinylestradiol) are degradable and are relatively hydrophobic, and in high-salinity waters reduced aqueous solubility promotes their adsorption onto suspended particles and sediments [84, 100–102]. Estrogens present in the wastewater may thus have been removed from the dissolved phase, which is the fraction sampled by the POCIS and analyzed in the bioassays, before the wastewater reached the sampling sites. The fate of estrogens in high-salinity reef waters nonetheless remains poorly understood [103], and this interpretation requires further research. The low ERα activity even at wastewater-impacted sites does not necessarily contradict the wastewater signal found in the target analysis, which is based on detected tracers (sucralose, fluconazole, lamotrigine) that are persistent, polar compounds that survive treatment and transport and reach the passive samplers, while not inducing ERα activity.

The PAH response, a proxy for industrial pollution like petroleum hydrocarbons [104], was generally low, with detectable responses at six out of nine tested sites, suggesting a widespread but low background presence of these compounds in the dissolved fraction sampled here. The widespread presence of these chemicals likely results from the long history of oil refining on the island of Curaçao [105], as well as intensified traffic both on land and at sea. PXR is activated by a wide and structurally diverse range of xenobiotics, including several pharmaceuticals and pesticides detected in this study (e.g., bezafibrate, diuron) [106]. Despite this, no PXR activity was detected in any of the samples, suggesting three possible explanations. First, the concentrations of PXR-activating compounds may have been present below the limit of detection of the bioassay (5.9 ng L^−1^ BEQ). Second, antagonistic interactions within the complex mixture may have suppressed activation: PXR binds multiple compounds simultaneously [107, 108], and one chemical can block another from activating the receptor [106]. Third, and most likely, the sampler itself is less amenable to the compounds that most strongly activate PXR and the aryl hydrocarbon receptor. Many of these (dioxin-like compounds, PAHs, PCBs) are hydrophobic, and the PES membrane used in the POCIS slows the uptake of compounds with a logK_ow_ > 2, and the HLB sorbent was selected for retention of polar to moderately polar compounds [49, 51, 52]. Previous effect-based studies applying passive sampling that recorded PXR and PAH activity did so in samplers that retain the more non-polar fraction [40, 50], which were not deployed here. Hence, the lack of PXR and PAH responses does not guarantee the absence of these compounds at ecotoxic levels on Curaçao’s coral reefs but does confirm low to negligible bioactivity for these endpoints in the water column at the studied sites. Similarly, the absence of cytotoxicity, which indicates acute cellular death, confirms low acute apical toxicity in the water column at the sampling sites [41].

Taken together, the effect-based methods showed that the time-integrated mixture of organic micropollutants was sufficiently bioactive to trigger a substantial cellular stress response (Nrf2) as well as low to moderate responses for endocrine (ERα and anti-AR) and metabolic (PAH) endpoints, but not potent enough to induce a PXR-mediated response or cytotoxicity. While the target chemical analysis pointed to the bays as the primary source of pollution, the effect-based assessment identified the highest biological activity (oxidative stress response) on the reefs. The CERs, the sum of the EBRQs, were predominantly driven by the Nrf2 responses and mirrored its pattern, with the highest overall risk found on the reefs. This demonstrates that target chemical analysis alone may not fully capture ecotoxicological risks and highlights the importance of integrating multiple lines of evidence in chemical water quality assessment.

### 3.3. Advantages and limitations of the applied methodology

The integrated approach of combining passive sampling with chemical analysis and EBMs provides a powerful framework for assessing water quality in dynamic coastal coral reef ecosystems. Passive sampling was coupled with passive flow monitors, providing a time-integrated mixture of chemical pollutants, capturing both chronic background levels and episodic pollution input, which can be missed by grab sampling. This passive sampling pre-concentrates pollutants on the sorbent over the deployment period, enabling sensitive detection of compounds at environmentally relevant concentrations that would be too low to measure directly in a water sample. This is particularly relevant in the often oligotrophic and strongly diluted waters overlying coral reefs.

Including bioassays is essential for understanding the potential ecotoxicological impact of these environmental chemical mixtures, as it avoids relying solely on the known toxicity of a few targeted compounds [39, 40]. Indeed, the compound-based and effect-based risk assessment approaches applied in the present study yielded complementary information on the mixture toxicity in Curaçao’s bays and coral reefs.

Despite its strengths, it is important to acknowledge the limitations of the applied methodology. First, the applied passive samplers (POCIS containing HLB sorbent) mainly extract the polar to moderately polar fraction of the micropollutants potentially present in the water column. Hydrophobic compounds, including many that may drive bioassay responses (e.g., PAH and PXR), are under-sampled, and their inclusion would require the parallel deployment of a non-polar sampler such as silicone rubber sheets [40, 49, 50], or the inclusion of sediment as an environmental compartment. Next, the accuracy of compound concentrations derived from passive samplers depends on the availability of compound-specific *Rs*. While *Rs* values were available for most detected compounds, they are often calibrated to freshwater conditions and under laboratory settings, introducing uncertainty when applied to dynamic tropical marine environments [35, 51]. This inherently leads to a probable misestimation of the actual time-weighted chemical water concentrations and thus of the true ecotoxicological risks. However, it can be argued that, despite this challenge, passive samplers still present a more realistic picture of actual chemical exposure than individual grab samples. Additionally, the compound-based chemical water quality assessment underestimates the calculated toxic pressures in two ways: i) the target chemical analysis inherently overlooks chemicals outside the target list as well as unknown chemicals, and ii) the availability of reliable toxicity data limits the number of chemicals that can be included in risk calculations. For example, msPAF values could be calculated only for the 14 detected compounds for which the necessary SSDs were available, and detected compounds such as DEET, desvenlafaxine, and naproxen, despite their presence and likely toxicity, did not contribute to the calculated mixture toxic pressure values. This highlights the shortcomings of relying solely on target analysis and reinforces the value of integrating untargeted approaches, such as non-target analysis and EBM, in chemical mixture risk assessment.

Furthermore, the ecotoxicological data used for the chemical risk assessments are predominantly derived from toxicity data for freshwater organisms, both for the SSDs underlying the msPAF calculation and those used for the derivation of the EBTs for the bioassays. Therefore, the specific sensitivities of marine organisms in general, and particularly for tropical coral reef organisms, are not directly represented. This warrants the application of a safety factor to account for unknown species sensitivities to toxicants as a precautionary approach, which is commonly applied when extrapolating toxicity data between species (groups), and was also applied here to derive interim EBTs for the marine environment. Similarly, a safety factor of 10 is also applied to derive the marine PNECs from the NORMAN DataBase, which were used in this study. Nonetheless, Yanagihara et al. [109] demonstrated that freshwater SSDs can be used to estimate saltwater SSDs reasonably well, suggesting that the applied msPAF approach has merit for estimating mixture toxicity even in coral reef environments for which few toxicity data are available.

Finally, a major challenge to identifying ecotoxicological effects on tropical marine biota is the lack of representative bioassays. There are only a few standardized bioassay protocols for marine organisms, and this is especially true for tropical coral reef species. Such protocols are under development [110, 111], and bioassays with reef-building organisms have shown their value in elucidating chemical pollution effects on coral reefs. For example, coral larvae suffered no acute mortality after direct exposure to local oil spill water from Curaçao in 2012, but showed reduced settlement, preventing them from developing into adult corals [112]. However, both acute and chronic effects of anthropogenic pollutants on coral reef organisms are at present understudied, and there is little insight into chemical pollution effects of coral reef structure and functioning. Hence, the marine ecotoxicological field would strongly benefit from the development of marine bioassays, targeted to local ecosystems, such as shallow-water tropical coral reefs.

### 3.4. Broader implications and management applications

#### 3.4.1. Broader implications

This study illustrates how land-based pollution reaches nearshore coral reefs: the inland bays of Curaçao act as reservoirs and sources of chemical contaminants, each with a distinct anthropogenic chemical fingerprint that reflects its local land use and pollution sources. In the dry season, the ranking of the bays by cumulative concentration follows the proportion of altered surface in their watersheds (Piscadera > Santa Martha > Awa Blanku; Table 1). The same spatial pattern was found independently for anthropogenic nitrogen: along the coast of Curaçao, macroalgal δ^15^N, a proxy for anthropogenic input, was elevated downstream of urbanized watersheds with a high proportion of altered surface [42, 43]. Nutrients and organic chemicals, as independent tracers, thus point to the same land-use-driven sources. Concentrations of most compounds were highest inside the bays and diluted towards the reefs. There was a strong seasonal dynamic, wherein rainfall during the wet season mobilizes pollutants from the watershed, elevating concentrations across all coastal waters, identifying the wet season as the critical period for contaminant delivery to the coastal zone, during which hydrological regimes retain water near the coast and may prolong exposure [45].

Critically, this seasonal pulse of pollution coincides with elevated sea surface temperatures and the reproductive and recruitment seasons of many reef-building corals and other marine organisms [30, 113]. Hence, the highest concentrations of land-based chemical stressors reach the reefs precisely when early life stages are most vulnerable to combined stress [26]. The effect-based assessment located the highest oxidative stress response at the reefs rather than inside the bays. Previously, macroalgal δ^15^N confirmed that land-derived material reaches the reef sites: elevated values occurred on the reefs themselves, most strongly at shallow depths and in the wet season [42]. Because the inland bays were not sampled in that study, δ^15^N cannot rank bays against reefs but it does show that terrestrial input is not confined to the bays. In this study, the POCIS were deployed at 9-11 m, below the depth range in which the δ^15^N signal was strongest, and the reef exposure measured here might therefore be a conservative estimate of the chemical exposure shallower reef communities experience. Together, these studies demonstrate that reef organisms are exposed to bioactive chemical mixtures at concentrations likely to induce ecotoxicological effects, which cannot be resolved by chemical analysis alone. Curaçao shares many characteristics with other Caribbean islands, including a growing (seasonal) population, increasing anthropogenic land use, reliance on tourism, and limited wastewater treatment infrastructure. Similar patterns of chemical pollution and associated ecotoxicological risks can therefore be expected across the region, suggesting the need for expanded monitoring and management attention.

#### 3.4.2. Management applications

Chemical pollution is one of several local stressors on the reefs of Curaçao, alongside fishing pressure, eutrophication, and the effects of climate change (e.g., ocean warming leading to bleaching). Addressing chemical pollution alone will not stop the decline of corals on reefs completely, but knowledge of the composition and effects of chemical mixtures allows interventions to be targeted at their sources, providing agency to local water managers and authorities to protect coral reefs, rather than relying on global actions to reduce climate change. The distinct fingerprint of each bay suggests that management responses should be tailored to local sources. Chronic urban pollution, such as the wastewater signature in Piscadera Bay, points toward infrastructure-focused interventions, i.e., upgrading wastewater treatment and improving sanitation systems to reduce continuous discharge of sewage-related contaminants [26, 114]. Episodic pollution mobilized during the wet season, from urban runoff and developed areas (e.g., hotels and residential development), could be addressed through watershed-focused interventions such as restoring natural buffer zones and coastal wetlands as filtration zones [26].

Conventional monitoring programs that focus on a limited list of compounds may not capture the cumulative toxicity of the full mixture to which reef organisms are exposed [60], as the presently found difference between chemical target- and effect-based assessments demonstrates. The multidisciplinary assessment used here, combining passive sampling with chemical analysis and bioassays, provides a framework for diagnosing pollution sources: a single indicator such as sucralose tracks wastewater inputs, while bioassays capture the integrated effects of the full mixture, including untargeted compounds. A simplified version combining key indicator chemicals with the most responsive bioassays could help guide and prioritize management actions in resource-limited regions, which is the de facto case for most tropical coral reef ecosystems.

#### 3.4.3 Future directions

A critical step in making this integrated approach applicable to coral reef ecosystems is to validate the applied methods for local conditions. Technically, in-situ calibration of passive samplers in tropical marine conditions would improve the accuracy of the derived water concentrations, and a parallel non-polar sampler would extend the assessment to the hydrophobic fraction. Bioassays using key reef organisms are being developed, but remain to be standardized for both individual chemicals and environmental mixtures. Such bioassays use the high sensitivity of early coral life stages with endpoints such as fertilization, larval survival, and settlement rates in response to specific contaminants like metals and UV filters [26, 115, 116] or to environmental mixtures [117]. Other sensitive reef invertebrates, like sea urchin embryotoxicity assays, measure developmental abnormalities [118], and the photosynthetic efficiency of algal symbionts indicates exposure to photosynthesis-inhibiting herbicides like diuron [26].

These studies still employ different methodologies and are rarely integrated with chemical analysis, which hinders the synthesis required for a robust, community-level risk model. Combining them to generate comparable species-specific data would provide the foundation for ecologically relevant toxicity thresholds for individual chemicals and mixtures and ultimately for reef-specific SSDs that remove much of the uncertainty of extrapolating from freshwater systems. Integrating these tools into monitoring frameworks would shift the nature of the work from documenting pollution to predicting and mitigating its ecological consequences, which is essential to preserve coral reef resilience against compounding chemical and climatic threats.

## List of abbreviations

ACN: acetonitrile
*anti*-AR: antagonism of the androgen receptor
BEQ: bioanalytical equivalent concentrations
CALUX: Chemically Activated LUciferase eXpression
CEC: contaminant of emerging concern
CER: cumulative ecotoxicological risk
C_W_: surface water concentrations
C_extr_: extract concentrations
DMSO: dimethyl sulfoxide
EBM: effect-based method
EBRQ: effect-based risk quotient
EBT: effect-based trigger
ER_α_: estrogen receptor alpha
LC-MS: liquid chromatography mass spectrometry
L_PFM_: relative PFM loss (see below)
MeOH: methanol
msPAF: multi-substance potentially affected fraction of species
Nrf2: nuclear factor erythroid 2-related factor 2
PFM: passive flow monitor
PNEC: predicted no-effect concentration
POCIS: polar organic chemical integrative sampler
PES: polyethersulfone
PXR: pregnane X receptor
REF: relative enrichment factor
R_extr_: extract-recovery
RQ: risk quotient
R_sorb_: sorbent-recovery
R_s_: sampling rate
SPE: solid-phase extraction
SSD: species sensitivity distribution
t: time

## Acknowledgements

This project received funding through the project “Land, Sea, and Society: Linking terrestrial pollutants and inputs to nearshore coral reef growth to identify novel conservation options for the Dutch Caribbean (SEALINK)” with project number NWOCA.2019.003 of the research program “Caribbean Research: a Multidisciplinary Approach” which is financed by the Dutch Research Council (NWO). Students Georgia Sarantidi, Stavriana Neokleous, and Vera van Schuilenburg are kindly acknowledged for assisting with sample collection. The Marine Park Rangers, Oscar, and the team at CARMABI are thanked for logistical support during fieldwork. We thank Sarit Kaserzon, Sara Ghorbani Gorji, and Chris Paxman from the University of Queensland for their methodological advice.

## CRediT author statement

**Nienke C.J. van de Loosdrecht**: Conceptualization, Methodology, Investigation, Formal Analysis, Data curation, Visualization, Writing – original draft. **Milo L. de Baat**: Conceptualization, Methodology, Investigation, Supervision, Writing – review & editing. **Mark J.A. Vermeij**: Conceptualization, Resources, Funding Acquisition, Writing – review & editing. **Patricia Aguilar-Alarcón**: Methodology, Investigation, Validation, Formal Analysis, Writing – review & editing. **Petra M. Visser:** Conceptualization, Methodology, Funding Acquisition, Investigation, Supervision, Writing – review & editing. **Jasper M. de Goeij:** Conceptualization, Methodology, Funding Acquisition, Investigation, Supervision, Project Administration, Writing – review & editing.

---

**Figure S1.**
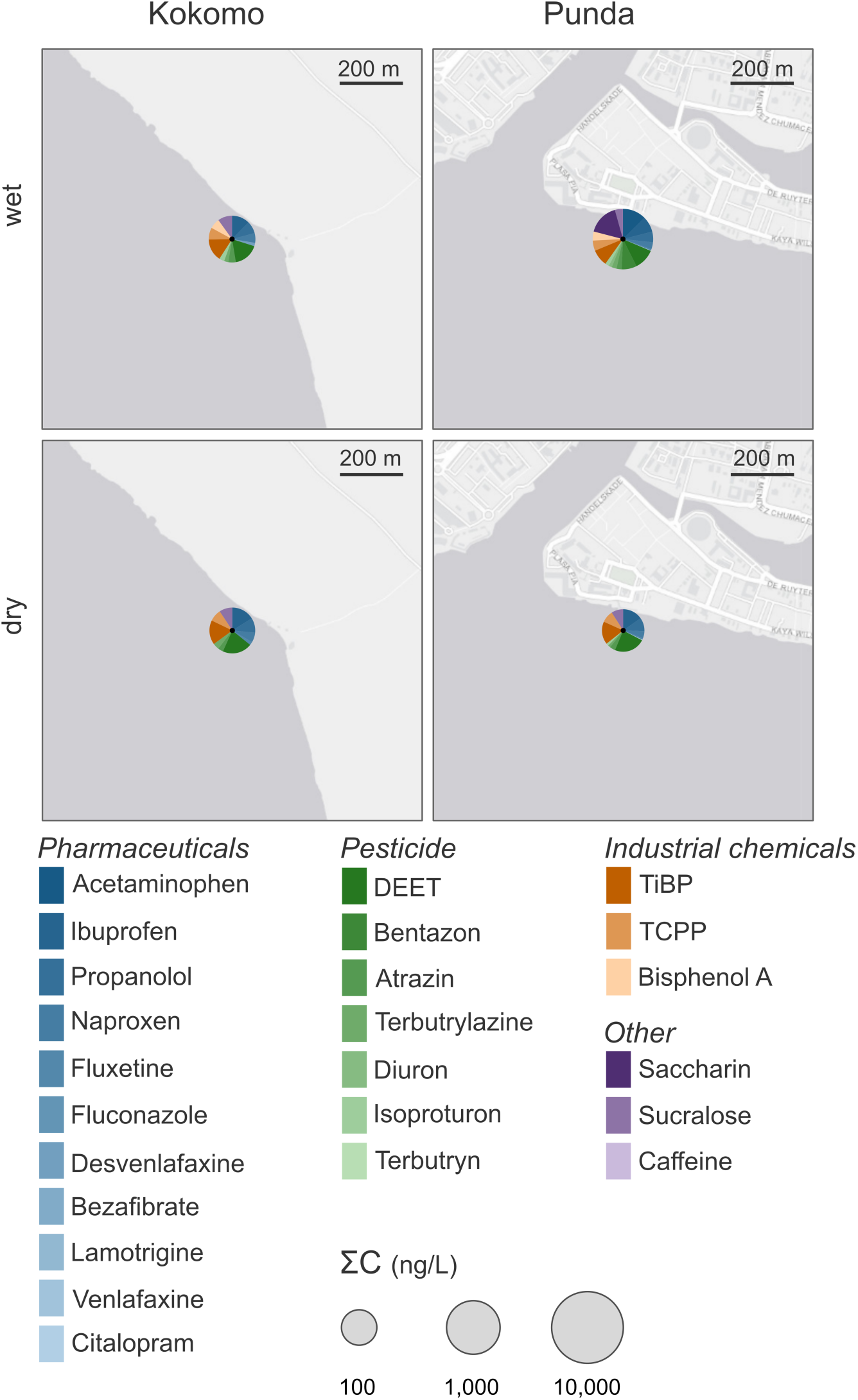
Spatial and seasonal concentrations of target compounds from passive sampler extracts of the coral reef sites Kokomo and Punda on Curaçao, southern Caribbean. Pie charts show the 24 quantified compounds collected by polar organic chemical integrative samplers (POCIS) and detected at each site during the wet (top) and dry (bottom) seasons of 2024. Pie size represents the total summed concentration of all compounds (ΣC, ng L^−1^, log scale), and slices show individual compounds (log_10_ (1+C)), colored by compound group.

## References

[1] Moberg F, Folke C, 1999. Ecological goods and services of coral reef ecosystems. Ecological Economics 29, 215–233. 10.1016/S0921-8009(99)00009-9

[2] Sobha TR, Vibija CP, Fahima P, 2023. Coral Reef: A Hot Spot of Marine Biodiversity. In: Sukumaran ST, T R K (eds) Conservation and Sustainable Utilization of Bioresources. Springer Nature Singapore, Singapore, pp 171–194

[3] Eddy TD, Lam VWY, Reygondeau G, Cisneros-Montemayor AM, Greer K, Palomares MLD, Bruno JF, Ota Y, Cheung WWL, 2021. Global decline in capacity of coral reefs to provide ecosystem services. One Earth 4, 1278–1285. 10.1016/j.oneear.2021.08.016

[4] Spalding M, Burke L, Wood SA, Ashpole J, Hutchison J, zu Ermgassen P, 2017. Mapping the global value and distribution of coral reef tourism. Marine Policy 82, 104–113. 10.1016/j.marpol.2017.05.014

[5] Hoegh-Guldberg O, Pendleton L, Kaup A, 2019. People and the changing nature of coral reefs. Regional Studies in Marine Science 30, 100699. 10.1016/j.rsma.2019.100699

[6] França FM, Benkwitt CE, Peralta G, Robinson JPW, Graham NAJ, Tylianakis JM, Berenguer E, Lees AC, Ferreira J, Louzada J, Barlow J, 2020. Climatic and local stressor interactions threaten tropical forests and coral reefs. Phil Trans R Soc B 375, 20190116. 10.1098/rstb.2019.0116

[7] Carlson RR, Foo SA, Asner GP, 2019. Land Use Impacts on Coral Reef Health: A Ridge-to-Reef Perspective. Front Mar Sci 6, 562. 10.3389/fmars.2019.00562

[8] Baum G, Januar HI, Ferse SCA, Kunzmann A, 2015. Local and Regional Impacts of Pollution on Coral Reefs along the Thousand Islands North of the Megacity Jakarta, Indonesia. PLOS ONE 10, e0138271. 10.1371/journal.pone.0138271

[9] Turak E, DeVantier L, Szava-Kovats R, Brodie J, 2021. Impacts of coastal land use change in the wet tropics on nearshore coral reefs: Case studies from Papua New Guinea. Marine Pollution Bulletin 168, 112445. 10.1016/j.marpolbul.2021.112445

[10] Fabricius KE, 2005. Effects of terrestrial runoff on the ecology of corals and coral reefs: review and synthesis. Marine Pollution Bulletin 50, 125–146. 10.1016/j.marpolbul.2004.11.028

[11] Strokal M, Bai Z, Franssen W, Hofstra N, Koelmans AA, Ludwig F, Ma L, van Puijenbroek P, Spanier JE, Vermeulen LC, van Vliet MTH, van Wijnen J, Kroeze C, 2021. Urbanization: an increasing source of multiple pollutants to rivers in the 21st century. npj Urban Sustainability 1, 24. 10.1038/s42949-021-00026-w

[12] Sheikh MA, Higuchi T, Fujimura H, Imo TS, Miyagi T, Oomori T, 2009. Contamination and impacts of new antifouling biocide Irgarol-1051 on subtropical coral reef waters. International Journal of Environmental Science & Technology 6, 353–358. 10.1007/BF03326073

[13] Risk MJ, Lapointe BE, Sherwood OA, Bedford BJ, 2009. The use of δ15N in assessing sewage stress on coral reefs. Marine Pollution Bulletin 58, 793–802. 10.1016/j.marpolbul.2009.02.008

[14] Häder D-P, Banaszak AT, Villafañe VE, Narvarte MA, González RA, Helbling EW, 2020. Anthropogenic pollution of aquatic ecosystems: Emerging problems with global implications. Science of The Total Environment 713, 136586. 10.1016/j.scitotenv.2020.136586

[15] Paytan A, Shellenbarger GG, Street JH, Gonneea ME, Davis K, Young MB, Moore WS, 2006. Submarine groundwater discharge: An important source of new inorganic nitrogen to coral reef ecosystems. Limnol Oceanogr 51, 343–348. 10.4319/lo.2006.51.1.0343

[16] D’Angelo C, Wiedenmann J, 2014. Impacts of nutrient enrichment on coral reefs: new perspectives and implications for coastal management and reef survival. Current Opinion in Environmental Sustainability 7, 82–93. 10.1016/j.cosust.2013.11.029

[17] Glasl B, Bourne DG, Frade PR, Thomas T, Schaffelke B, Webster NS, 2019. Microbial indicators of environmental perturbations in coral reef ecosystems. Microbiome 7, 94. 10.1186/s40168-019-0705-7

[18] van Dam JW, Negri AP, Uthicke S, Mueller JF, 2011. Chemical Pollution on Coral Reefs: Exposure and Ecological Effects

[19] Khan S, Naushad Mu, Govarthanan M, Iqbal J, Alfadul SM, 2022. Emerging contaminants of high concern for the environment: Current trends and future research. Environmental Research 207, 112609. 10.1016/j.envres.2021.112609

[20] Nishmitha PS, Akhilghosh KA, Aiswriya VP, Ramesh A, Muthuchamy M, Muthukumar A, 2025. Understanding emerging contaminants in water and wastewater: A comprehensive review on detection, impacts, and solutions. Journal of Hazardous Materials Advances 18, 100755. 10.1016/j.hazadv.2025.100755

[21] Puri M, Gandhi K, Kumar MS, 2023. Emerging environmental contaminants: A global perspective on policies and regulations. Journal of Environmental Management 332, 117344. 10.1016/j.jenvman.2023.117344

[22] Yadav D, Rangabhashiyam S, Verma P, Singh P, Devi P, Kumar P, Hussain CM, Gaurav GK, Kumar KS, 2021. Environmental and health impacts of contaminants of emerging concerns: Recent treatment challenges and approaches. Chemosphere 272, 129492. 10.1016/j.chemosphere.2020.129492

[23] Ng B, Quinete N, Maldonado S, Lugo K, Purrinos J, Briceño H, Gardinali P, 2021. Understanding the occurrence and distribution of emerging pollutants and endocrine disruptors in sensitive coastal South Florida Ecosystems. Science of The Total Environment 757, 143720. 10.1016/j.scitotenv.2020.143720

[24] Wear SL, Thurber RV, 2015. Sewage pollution: mitigation is key for coral reef stewardship. Annals of the New York Academy of Sciences 1355, 15–30. 10.1111/nyas.12785

[25] Krzeminski P, Tomei MC, Karaolia P, Langenhoff A, Almeida CMR, Felis E, Gritten F, Andersen HR, Fernandes T, Manaia CM, 2019. Performance of secondary wastewater treatment methods for the removal of contaminants of emerging concern implicated in crop uptake and antibiotic resistance spread: A review. Science of the Total Environment 648, 1052–1081

[26] Nalley EM, Tuttle LJ, Barkman AL, Conklin EE, Wulstein DM, Richmond RH, Donahue MJ, 2021. Water quality thresholds for coastal contaminant impacts on corals: A systematic review and meta-analysis. Science of The Total Environment 794, 148632. 10.1016/j.scitotenv.2021.148632

[27] Bainbridge Z, Lewis S, Bartley R, Fabricius K, Collier C, Waterhouse J, Garzon-Garcia A, Robson B, Burton J, Wenger A, Brodie J, 2018. Fine sediment and particulate organic matter: A review and case study on ridge-to-reef transport, transformations, fates, and impacts on marine ecosystems. Marine Pollution Bulletin 135, 1205–1220. 10.1016/j.marpolbul.2018.08.002

[28] Govers LL, Lamers LPM, Bouma TJ, Eygensteyn J, De Brouwer JHF, Hendriks AJ, Huijbers CM, Van Katwijk MM, 2014. Seagrasses as indicators for coastal trace metal pollution: A global meta-analysis serving as a benchmark, and a Caribbean case study. Environmental Pollution 195, 210–217. 10.1016/j.envpol.2014.08.028

[29] Page CE, Ainsworth TD, Leggat W, Egan S, Gupta AS, Raoult V, Gaston TF, 2023. Localising terrestrially derived pollution inputs to threatened near-shore coral reefs through stable isotope, water quality and oceanographic analysis. Marine Pollution Bulletin 193, 115193. 10.1016/j.marpolbul.2023.115193

[30] De Jong C, Van Os I, Sepúlveda-Rodríguez G, De Baat ML, Schoepf V, 2025. High-resolution temporal assessment of physicochemical variability and water quality in tropical semi-enclosed bays and coral reefs. Science of The Total Environment 968, 178810. 10.1016/j.scitotenv.2025.178810

[31] Boonstra H, de Baat ML, van der Meer F, Besselink H, Roessink I, Kraak MHS, 2025. Capturing temporal variation in aquatic ecotoxicological risks: Chemical- versus effect-based assessment. Science of The Total Environment 967, 178797. 10.1016/j.scitotenv.2025.178797

[32] Brodie J, Lewis S, Collier C, Wooldridge S, Bainbridge ZT, Waterhouse J, Rasheed M, Honchin C, Holmes G, Fabricius K, 2017. Setting ecologically relevant targets for river pollutant loads to meet marine water quality requirements for the Great Barrier Reef, Australia: A preliminary methodology and analysis. Ocean & Coastal Management 143, 136–147. 10.1016/J.OCECOAMAN.2016.09.028

[33] Den Haan J, Huisman J, Brocke HJ, Goehlich H, Latijnhouwers KRW, Van Heeringen S, Honcoop SAS, Bleyenberg TE, Schouten S, Cerli C, Hoitinga L, Vermeij MJA, Visser PM, 2016. Nitrogen and phosphorus uptake rates of different species from a coral reef community after a nutrient pulse. Sci Rep 6, 28821. 10.1038/srep28821

[34] Vrana B, Allan IJ, Greenwood R, Mills GA, Dominiak E, Svensson K, Knutsson J, Morrison G, 2005. Passive sampling techniques for monitoring pollutants in water. TrAC Trends in Analytical Chemistry 24, 845–868. 10.1016/j.trac.2005.06.006

[35] MacKeown H, Benedetti B, Di Carro M, Magi E, 2022. The study of polar emerging contaminants in seawater by passive sampling: A review. Chemosphere 299, 134448. 10.1016/j.chemosphere.2022.134448

[36] Sigmund G, Ågerstrand M, Antonelli A, Backhaus T, Brodin T, Diamond ML, Erdelen WR, Evers DC, Hofmann T, Hueffer T, Lai A, Torres JPM, Mueller L, Perrigo AL, Rillig MC, Schaeffer A, Scheringer M, Schirmer K, Tlili A, Soehl A, Triebskorn R, Vlahos P, vom Berg C, Wang Z, Groh KJ, 2023. Addressing chemical pollution in biodiversity research. Global Change Biology 29, 3240–3255. 10.1111/gcb.16689

[37] Markert N, Rhiem S, Trimborn M, Guhl B, 2020. Mixture toxicity in the Erft River: assessment of ecological risks and toxicity drivers. Environmental Sciences Europe 32, 51. 10.1186/s12302-020-00326-5

[38] Weisner O, Frische T, Liebmann L, Reemtsma T, Roß-Nickoll M, Schäfer RB, Schäffer A, Scholz-Starke B, Vormeier P, Knillmann S, Liess M, 2021. Risk from pesticide mixtures – The gap between risk assessment and reality. Science of The Total Environment 796, 149017. 10.1016/j.scitotenv.2021.149017

[39] Neale PA, Escher BI, de Baat ML, Dechesne M, Dingemans MML, Enault J, Pronk GJ, Smeets PWMH, Leusch FDL, 2023. Application of Effect-Based Methods to Water Quality Monitoring: Answering Frequently Asked Questions by Water Quality Managers, Regulators, and Policy Makers. Environ Sci Technol 57, 6023–6032. 10.1021/acs.est.2c06365

[40] De Baat ML, Van der Oost R, Van der Lee GH, Wieringa N, Hamers T, Verdonschot PFM, De Voogt P, Kraak MHS, 2020. Advancements in effect-based surface water quality assessment. Water Research 183, 116017. 10.1016/j.watres.2020.116017

[41] Escher BI, Neale PA, 2021. Effect-Based Trigger Values for Mixtures of Chemicals in Surface Water Detected with In Vitro Bioassays. Environmental Toxicology and Chemistry 40, 487–499. 10.1002/etc.4944

[42] van de Loosdrecht NCJ, Visser PM, de Baat ML, Vermeij MJA, de Goeij JM, 2026. Macroalgal δ15N indicates land-derived elemental pollution on nearshore coral reefs of Curaçao, southern Caribbean. Marine Pollution Bulletin 223, 118967. 10.1016/j.marpolbul.2025.118967

[43] Sandin SA, Alcantar E, Clark R, De León R, Dilrosun F, Edwards CB, Estep AJ, Eynaud Y, French BJ, Fox MD, Grenda D, Hamilton SL, Kramp H, Marhaver KL, Miller SD, Roach TNF, Seferina G, Silveira CB, Smith JE, Zgliczynski BJ, Vermeij MJA, 2022. Benthic assemblages are more predictable than fish assemblages at an island scale. Coral Reefs 41, 1031–1043. 10.1007/s00338-022-02272-5

[44] Sanchez Barranco V, Schellenberg L, Mienis F, Brussaard CPD, Haas AF, de Nooijer LJ, 2025. Seasonal changes in bay water column properties and their influence on the distribution of dissolved and particulate substances along the south coast of Curaçao (Caribbean Sea). Marine Pollution Bulletin 212, 117545. 10.1016/j.marpolbul.2025.117545

[45] Bertoncelj V, Mienis F, Stocchi P, van Sebille E, 2025. Flow patterns, hotspots, and connectivity of land-derived substances at the sea surface of Curaçao in the southern Caribbean. Ocean Science 21, 945–964. 10.5194/os-21-945-2025

[46] Curaçao CB of S, 2020. Population. Central Bureau of Statistics Curaçao. https://www.cbs.cw/population. Accessed 3 June 2026

[47] Curaçao CB of S, 2025. Tourism. Central Bureau of Statistics Curaçao. https://www.cbs.cw/tourism-2. Accessed 3 June 2026

[48] Meteorological Department Curaçao

[49] de Baat ML, Narain-Ford DM, de Weert J, Giesen D, Beeltje H, Hamers T, Helmus R, de Voogt P, Kraak MHS, 2024. Passive sampler housing and sorbent type determine aquatic micropollutant adsorption and subsequent bioassay responses. Environmental Pollution 357, 124488. 10.1016/j.envpol.2024.124488

[50] de Weert J, Smedes F, Beeltje H, de Zwart D, Hamers T, 2020. Time integrative sampling properties of Speedisk and silicone rubber passive samplers determined by chemical analysis and in vitro bioassay testing. Chemosphere 259, 127498. 10.1016/j.chemosphere.2020.127498

[51] Harman C, Allan I, Thomas KV, 2012. 1.14 - Passive Sampling of Organic Contaminants in Waters. In: Pawliszyn J (ed) Comprehensive Sampling and Sample Preparation. Academic Press, Oxford, pp 265–280

[52] Vermeirssen ELM, Dietschweiler C, Escher BI, van der Voet J, Hollender J, 2012. Transfer Kinetics of Polar Organic Compounds over Polyethersulfone Membranes in the Passive Samplers Pocis and Chemcatcher. Environmental Science & Technology 46, 6759–6766. 10.1021/es3007854

[53] Togola A, Budzinski H, 2007. Development of Polar Organic Integrative Samplers for Analysis of Pharmaceuticals in Aquatic Systems. Analytical Chemistry 79, 6734–6741. 10.1021/ac070559i

[54] Charlestra L, Amirbahman A, Courtemanch DL, Alvarez DA, Patterson H, 2012. Estimating pesticide sampling rates by the polar organic chemical integrative sampler (POCIS) in the presence of natural organic matter and varying hydrodynamic conditions. Environmental Pollution 169, 98–104. 10.1016/j.envpol.2012.05.001

[55] Alvarez DA, Petty JD, Huckins JN, Jones- Lepp TL, Getting DT, Goddard JP, Manahan SE, 2004. Development of a passive, in situ, integrative sampler for hydrophilic organic contaminants in aquatic environments. Environmental Toxicology and Chemistry 23, 1640–1648. 10.1897/03-603

[56] Godlewska K, Stepnowski P, Paszkiewicz M, 2021. Pollutant analysis using passive samplers: principles, sorbents, calibration and applications. A review. Environmental Chemistry Letters 19, 465–520. 10.1007/s10311-020-01079-6

[57] O’Brien D, Bartkow M, Mueller JF, 2011. Determination of deployment specific chemical uptake rates for SDB-RPD Empore disk using a passive flow monitor (PFM). Chemosphere 83, 1290–1295. 10.1016/j.chemosphere.2011.02.089

[58] Specker JC, Aguilar-Alarcón P, Helmus R, Felzel E, Besseling H, Behnisch P, de Baat ML, Praetorius A, Sutton NB, van Wezel AP, 2026. Beyond Targets: Benefits of a Multi-Assessment Approach in Determining Chemical Water Quality for Water Reuse

[59] van der Linden SC, von Bergh ARM, van Vught-Lussenburg BMA, Jonker LRA, Teunis M, Krul CAM, van der Burg B, 2014. Development of a panel of high-throughput reporter-gene assays to detect genotoxicity and oxidative stress. Mutation Research/Genetic Toxicology and Environmental Mutagenesis 760, 23–32. 10.1016/j.mrgentox.2013.09.009

[60] Escher BI, van Daele C, Dutt M, Tang JYM, Altenburger R, 2013. Most Oxidative Stress Response In Water Samples Comes From Unknown Chemicals: The Need For Effect-Based Water Quality Trigger Values. Environ Sci Technol 47, 7002–7011. 10.1021/es304793h

[61] NORMAN Network. NORMAN Ecotoxicology Database - Lowest PNECs

[62] Dulio V, van Bavel B, Brorström-Lundén E, Harmsen J, Hollender J, Schlabach M, Slobodnik J, Thomas K, Koschorreck J, 2018. Emerging pollutants in the EU: 10 years of NORMAN in support of environmental policies and regulations. Environmental Sciences Europe 30, 5. 10.1186/s12302-018-0135-3

[63] Ouédraogo D-Y, Mell H, Perceval O, Burga K, Domart-Coulon I, Hédouin L, Delaunay M, Guillaume MMM, Castelin M, Calvayrac C, Kerkhof O, Sordello R, Reyjol Y, Ferrier-Pagès C, 2023. What are the toxicity thresholds of chemical pollutants for tropical reef-building corals? A systematic review. Environ Evid 12, 4. 10.1186/s13750-023-00298-y

[64] Ouédraogo D-Y, Sordello R, Brugneaux S, Burga K, Calvayrac C, Castelin M, Domart-Coulon I, Ferrier-Pagès C, Guillaume MMM, Hédouin L, Joannot P, Perceval O, Reyjol Y, 2020. What evidence exists on the impacts of chemicals arising from human activity on tropical reef-building corals? A systematic map protocol. Environ Evid 9, 18. 10.1186/s13750-020-00203-x

[65] de Zwart D, Posthuma L, 2005. Complex mixture toxicity for single and multiple species: Proposed methodologies. Environmental Toxicology and Chemistry 24, 2665–2676. 10.1897/04-639R.1

[66] Klepper O, Bakker J, Traas TP, van de Meent D, 1998. Mapping the potentially affected fraction (PAF) of species as a basis for comparison of ecotoxicological risks between substances and regions. Journal of Hazardous Materials 61, 337–344

[67] Posthuma L, van Gils J, Zijp MC, Van De Meent DIK, de Zwart D, 2019. Species sensitivity distributions for use in environmental protection, assessment, and management of aquatic ecosystems for 12 386 chemicals. Environmental Toxicology and Chemistry 38, 905–917

[68] Kwok KW, Leung KM, Lui GS, Chu VK, Lam PK, Morritt D, Maltby L, Brock TC, Van den Brink PJ, Warne MSJ, Crane M, 2007. Comparison of tropical and temperate freshwater animal species’ acute sensitivities to chemicals: Implications for deriving safe extrapolation factors. Integrated Environmental Assessment and Management 3, 49–67. 10.1002/ieam.5630030105

[69] R. Core Team, 2024. R: A language and environment for statistical computing. Vienna, Austria: R Foundation for Statistical Computing. https://www.R-project.org/

[70] Posit Team, 2024. RStudio: integrated development environment for R. Posit Software, PBC. http://www.posit.co/

[71] Oksanen J, Simpson GL, Blanchet FG, Kindt R, Legendre P, Minchin PR, O’hara RB, Solymos P, Stevens MHH, Szoecs E, 2001. Vegan: community ecology package. (No Title)

[72] Anderson MJ, 2001. A new method for non-parametric multivariate analysis of variance. Austral Ecology 26, 32–46. 10.1111/j.1442-9993.2001.01070.pp.x

[73] Anderson MJ, 2006. Distance-Based Tests for Homogeneity of Multivariate Dispersions. Biometrics 62, 245–253. 10.1111/j.1541-0420.2005.00440.x

[74] Benjamini Y, Hochberg Y, 1995. Controlling the False Discovery Rate: A Practical and Powerful Approach to Multiple Testing. Journal of the Royal Statistical Society: Series B (Methodological) 57, 289–300. 10.1111/j.2517-6161.1995.tb02031.x

[75] Lowe RJ, Falter JL, 2015. Oceanic Forcing of Coral Reefs. Annual Review of Marine Science 7:43–66

[76] Snijders K, van Breukelen B, Langeveld JG, Wit M, 2023. Urban Wastewater Management of Willemstad, Curaçao, and Environmental Implications: A Qualitative and Quantitative Investigation on Urban Wastewater Fluxes. Delft University of Technology, Faculty of Civil Engineering & Geosciences https://resolv er tudelftnl/uuid: 9112850b-9f17-4c4b-8f57-9db54c517e88

[77] Van Duyl FC, Post VEA, Van Breukelen BM, Bense V, Visser PM, Meesters EH, Koeniger P, Vermeij MJA, 2024. Composition and distribution of the near-shore waters bordering the coral reefs of Aruba, Bonaire, and Curaçao in the Southern Caribbean. Marine Pollution Bulletin 209, 117297. 10.1016/j.marpolbul.2024.117297

[78] Li J, Li X, Li Y, Liu H, Wang Q, 2025. Artificial sweeteners in wastewater treatment plants: A systematic review of global occurrence, distribution, removal, and degradation pathways. Journal of Hazardous Materials 494, 138644. 10.1016/j.jhazmat.2025.138644

[79] Singh SP, Azua A, Chaudhary A, Khan S, Willett KL, Gardinali PR, 2010. Occurrence and distribution of steroids, hormones and selected pharmaceuticals in South Florida coastal environments. Ecotoxicology 19, 338–350. 10.1007/s10646-009-0416-0

[80] Jones R, Kerswell A, 2003. Phytotoxicity of Photosystem II (PSII) herbicides to coral. Mar Ecol Prog Ser 261, 149–159. 10.3354/meps261149

[81] Finn J, Hui M, Li V, Lorenzi V, de la Paz N, Cheng SH, Lai-Chan L, Schlenk D, 2012. Effects of propranolol on heart rate and development in Japanese medaka (Oryzias latipes) and zebrafish (Danio rerio). Aquatic Toxicology 122–123, 214–221. 10.1016/j.aquatox.2012.06.013

[82] Melefa TD, Andong FA, Hinmikaiye FF, Ozota EE, Nwoko M-C, Ubah VCS, Ugwu GN, Nwani CD, 2025. Psychoactive drug propranolol modulates behavioral, acetylcholinesterase, and oxidative stress parameters in freshwater African Sharptooth Catfish Clarias gariepinus. Journal of Aquatic Animal Health 37, 66–74. 10.1093/jahafs/vsaf004

[83] Mathias FT, Fockink DH, Disner GR, Prodocimo V, Ribas JLC, Ramos LP, Cestari MM, Silva de Assis HC, 2018. Effects of low concentrations of ibuprofen on freshwater fish Rhamdia quelen. Environmental Toxicology and Pharmacology 59, 105–113. 10.1016/j.etap.2018.03.008

[84] Zhang J, Yang G-P, Li Q, Cao X, Liu G, 2013. Study on the sorption behaviour of estrone on marine sediments. Marine Pollution Bulletin 76, 220–226. 10.1016/j.marpolbul.2013.08.038

[85] Wang Z, Kwok KWH, Lui GCS, Zhou G-J, Lee J-S, Lam MHW, Leung KMY, 2014. The difference between temperate and tropical saltwater species’ acute sensitivity to chemicals is relatively small. Chemosphere 105, 31–43. 10.1016/j.chemosphere.2013.10.066

[86] Zimdahl RL, 2018. Chapter 16 - Properties and Uses of Herbicides. In: Fundamentals of Weed Science (Fifth Edition). Academic Press, pp 463–499

[87] Benedetti M, Giuliani ME, Mezzelani M, Nardi A, Pittura L, Gorbi S, Regoli F, 2022. Emerging environmental stressors and oxidative pathways in marine organisms: Current knowledge on regulation mechanisms and functional effects. Biocell 46, 37

[88] Reiter EB, Escher BI, Siebert U, Jahnke A, 2022. Activation of the xenobiotic metabolism and oxidative stress response by mixtures of organic pollutants extracted with in-tissue passive sampling from liver, kidney, brain and blubber of marine mammals. Environment International 165, 107337. 10.1016/j.envint.2022.107337

[89] Silva SB, Gregório KP, de Freitas RMP, Destro ALF, Salaro AL, Zuanon JAS, Gonçalves RV, Freitas MB, 2023. EXPOSURE TO THE HERBICIDE ATRAZINE INDUCES OXIDATIVE IMBALANCE, MORPHOLOGICAL DAMAGE AND DECREASED SURVIVAL IN JUVENILE FISH. Bioscience Journal 39

[90] Duarte IA, Reis-Santos P, Novais SC, Rato LD, Lemos MFL, Freitas A, Pouca ASV, Barbosa J, Cabral HN, Fonseca VF, 2020. Depressed, hypertense and sore: Long-term effects of fluoxetine, propranolol and diclofenac exposure in a top predator fish. Science of The Total Environment 712, 136564. 10.1016/j.scitotenv.2020.136564

[91] Parolini M, Ghilardi A, De Felice B, Del Giacco L, 2019. Environmental concentration of fluoxetine disturbs larvae behavior and increases the defense response at molecular level in zebrafish (Danio rerio). Environmental Science and Pollution Research 26, 34943–34952. 10.1007/s11356-019-06619-4

[92] Feijão E, Cruz de Carvalho R, Duarte IA, Matos AR, Cabrita MT, Utkin AB, Caçador I, Marques JC, Novais SC, Lemos MF, 2022. Fluoxetine induces photochemistry-derived oxidative stress on Ulva lactuca. Frontiers in Environmental Science 10, 963537

[93] Chatzigianni M, Pavlou P, Siamidi A, Vlachou M, Varvaresou A, Papageorgiou S, 2022. Environmental impacts due to the use of sunscreen products: a mini-review. Ecotoxicology 31, 1331–1345. 10.1007/s10646-022-02592-w

[94] de Jager TL, Cockrell AE, Du Plessis SS, 2017. Ultraviolet Light Induced Generation of Reactive Oxygen Species. In: Ahmad SI (ed) Ultraviolet Light in Human Health, Diseases and Environment. Springer International Publishing, Cham, pp 15–23

[95] Rasher DB, Hay ME, 2010. Chemically rich seaweeds poison corals when not controlled by herbivores. Proceedings of the National Academy of Sciences 107, 9683–9688. 10.1073/pnas.0912095107

[96] Reigel AM, Easson CG, Fiore CL, Apprill A, 2024. Sponge exhalent metabolites influence coral reef picoplankton dynamics. Scientific Reports 14, 31394. 10.1038/s41598-024-82995-3

[97] Houtman CJ, Brewster K, ten Broek R, Duijve B, van Oorschot Y, Rosielle M, Lamoree MH, Steen RJCA, 2021. Characterisation of (anti-)progestogenic and (anti-)androgenic activities in surface and wastewater using high resolution effectdirected analysis. Environment International 153, 106536. 10.1016/j.envint.2021.106536

[98] U.S. Environmental Protection Agency. CompTox Chemicals Dashboard 2.5.x - DEET, 134–62-3 - DTXSID2021995. CompTox Chemicals Dashboard. https://comptox.epa.gov/dashboard/chemical/invitrodb/DTXSID2021995

[99] Välitalo P, Perkola N, Seiler T-B, Sillanpää M, Kuckelkorn J, Mikola A, Hollert H, Schultz E, 2016. Estrogenic activity in Finnish municipal wastewater effluents. Water Research 88, 740–749. 10.1016/j.watres.2015.10.056

[100] do Nascimento MTL, Santos AD de O, Felix LC, Gomes G, de Oliveira e Sá M, da Cunha DL, Vieira N, Hauser-Davis RA, Baptista Neto JA, Bila DM, 2018. Determination of water quality, toxicity and estrogenic activity in a nearshore marine environment in Rio de Janeiro, Southeastern Brazil. Ecotoxicology and Environmental Safety 149, 197–202. 10.1016/j.ecoenv.2017.11.045

[101] Bowman JC, Zhou JL, Readman JW, 2002. Sediment–water interactions of natural oestrogens under estuarine conditions. Marine Chemistry 77, 263–276. 10.1016/S0304-4203(02)00006-3

[102] Faber M, James CA, Harding LB, da Silva DA, Sofield RM, 2025. Estrogenic activity of mixtures in the Salish Sea: The use of high throughput toxicity data with chemical information from fish bile and other matrices. Plos one 20, epone. 0323865

[103] Bosker T, Santoro G, Melvin SD, 2017. Salinity and sensitivity to endocrine disrupting chemicals: A comparison of reproductive endpoints in small-bodied fish exposed under different salinities. Chemosphere 183, 186–196. 10.1016/j.chemosphere.2017.05.063

[104] Neff JM, Stout SA, Gunster DG, 2005. Ecological risk assessment of polycyclic aromatic hydrocarbons in sediments: Identifying sources and ecological hazard. Integrated Environmental Assessment and Management 1, 22–33. 10.1897/IEAM_2004a-016.1

[105] Bond D, 2017. Oil in the Caribbean: Refineries, Mangroves, and the Negative Ecologies of Crude Oil. Comp Stud Soc Hist 59, 600–628. 10.1017/S0010417517000184

[106] Creusot N, Garoche C, Grimaldi M, Boulahtouf A, Chiavarina B, Bourguet W, Balaguer P, 2021. A Comparative Study of Human and Zebrafish Pregnane X Receptor Activities of Pesticides and Steroids Using In Vitro Reporter Gene Assays. Frontiers in Endocrinology Volume 12–2021

[107] Delfosse V, Dendele B, Huet T, Grimaldi M, Boulahtouf A, Gerbal-Chaloin S, Beucher B, Roecklin D, Muller C, Rahmani R, Cavaillès V, Daujat-Chavanieu M, Vivat V, Pascussi J-M, Balaguer P, Bourguet W, 2015. Synergistic activation of human pregnane X receptor by binary cocktails of pharmaceutical and environmental compounds. Nature Communications 6, 8089. 10.1038/ncomms9089

[108] de Sousa G, Nawaz A, Cravedi J-P, Rahmani R, 2014. A Concentration Addition Model to Assess Activation of the Pregnane X Receptor (PXR) by Pesticide Mixtures Found in the French Diet. Toxicological Sciences 141, 234–243. 10.1093/toxsci/kfu120

[109] Yanagihara M, Hiki K, Iwasaki Y, 2022. Can Chemical Toxicity in Saltwater Be Predicted from Toxicity in Freshwater? A Comprehensive Evaluation Using Species Sensitivity Distributions. Environmental Toxicology and Chemistry 41, 2021–2027. 10.1002/etc.5354

[110] Davey JL, Finlayson KA, Limpus CJ, van de Merwe JP, 2026. Species-specific bioassays assess toxicity in sympatric marine turtles. Regional Studies in Marine Science 102, 105441. 10.1016/j.rsma.2026.105441

[111] Brefeld D, Di Mauro V, Kellermann MY, Nietzer S, Moeller M, Lütjens LH, Pawlowski S, Petersen-Thiery M, Schupp PJ, 2024. Acute Toxicity Assays with Adult Coral Fragments: A Method for Standardization. Toxics 12, 1. 10.3390/toxics12010001

[112] Hartmann AC, Sandin SA, Chamberland VF, Marhaver KL, De Goeij JM, Vermeij MJ, 2015. Crude oil contamination interrupts settlement of coral larvae after direct exposure ends. Marine Ecology Progress Series 536, 163–173

[113] Vermeij MJA, Sampayo E, Bröker K, Bak RPM, 2004. The reproductive biology of closely related coral species: gametogenesis in Madracis from the southern Caribbean. Coral Reefs 23, 206–214. 10.1007/s00338-004-0368-z

[114] Zhao H, Yuan M, Strokal M, Wu HC, Liu X, Murk A, Kroeze C, Osinga R, 2021. Impacts of nitrogen pollution on corals in the context of global climate change and potential strategies to conserve coral reefs. Science of The Total Environment 774, 145017. 10.1016/j.scitotenv.2021.145017

[115] Miller IB, Moeller M, Kellermann MY, Nietzer S, Di Mauro V, Kamyab E, Pawlowski S, Petersen-Thiery M, Schupp PJ, 2022. Towards the development of standardized bioassays for corals: Acute toxicity of the UV filter benzophenone-3 to scleractinian coral larvae. Toxics 10, 244

[116] Nordborg FM, Brinkman DL, Ricardo GF, Agustí S, Negri AP, 2021. Comparative sensitivity of the early life stages of a coral to heavy fuel oil and UV radiation. Science of The Total Environment 781, 146676. 10.1016/j.scitotenv.2021.146676

[117] Latijnhouwers KRW, Van der Last AJ, Virdis F, Le Trocquer N, Tuijten S, Chamberland VF, Miller MW, 2026. Effects-based assessment of water and benthic habitat quality using coral larvae. Coral Reefs. 10.1007/s00338-026-02855-6

[118] Perina FC, de Souza Abessa DM, Pinho GLL, Fillmann G, 2011. Comparative toxicity of antifouling compounds on the development of sea urchin. Ecotoxicology 20, 1870–1880. 10.1007/s10646-011-0725-y

